# Host senescence and urbanized conditions drive bacterial communities of the freshwater sponge *Spongilla lacustris*

**DOI:** 10.64898/2026.01.07.698122

**Authors:** Chloën Lhomme, Benoit Paix, Nicole J. de Voogd

## Abstract

Freshwater sponge holobionts remain poorly characterized despite their ecological importance and exposure to strong anthropogenic pressures. Here, we investigated spatio-temporal dynamics of the bacterial microbiome within the freshwater sponge *Spongilla lacustris* across three sites with contrasting nutrient levels in the urban and peri-urban canal systems of Leiden (The Netherlands). Sponge and freshwater samples were collected every two weeks from October to December 2021, covering major stages of the adult sponge life cycle, including gemmulation, bleaching, and senescence. Using 16S rRNA gene metabarcoding, major differences in the sponge microbiome diversity were observed through a temporal dynamic, being associated with host physiological changes. Senescent and bleached sponges were characterized by an increased β-diversity dispersion, consistent with a potential dysbiosis process following the Anna Karenina Principle. A wide range of biomarker taxa were associated with this bleached stage and may correspond to opportunistic microbial scavengers degrading organic matter released during host decomposition. To a lesser extent, spatial patterns were also observed: ammonium- and phosphate-enriched sites harbored a limited number of discriminant taxa, characterised primarily as denitrifiers and methanotrophs, suggesting nutrient-driven functional structuring. Together, these results indicate that *S. lacustris* microbiomes are primarily shaped by host senescence, while environmental eutrophication influences specific taxa, providing new insights into freshwater sponge holobiont ecology under urbanized conditions.

**Graphical abstract:** 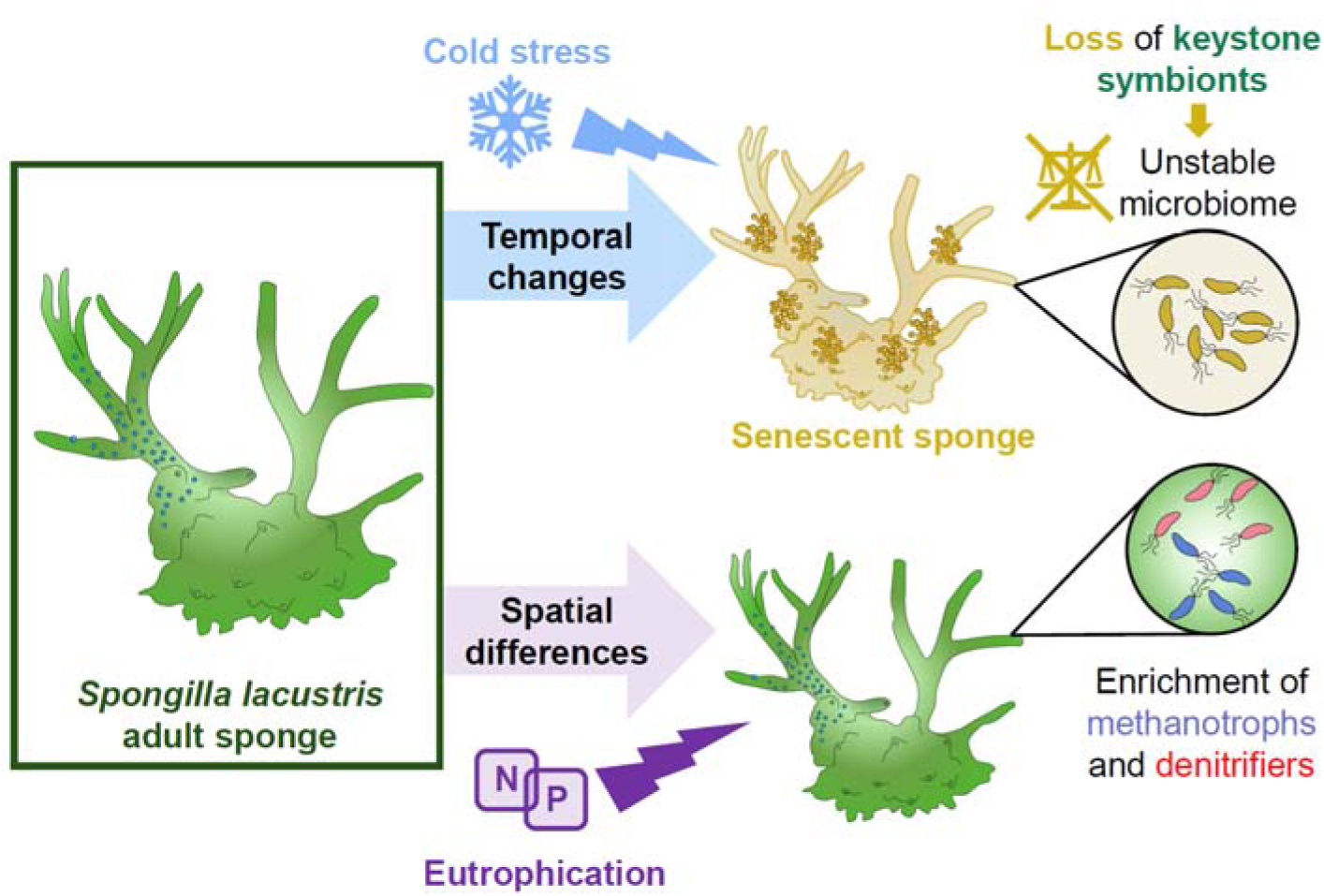

## Introduction

As ecosystem engineers, sponges provide key ecological roles within their environments. Among others, they contribute to the cycling of dissolved organic matter through their filter-feeding activities (Rix *et al*., 2017; Ribes *et al*., 2023), and act as a stable habitat for a large diversity of fauna (such as crustaceans, mollusks, and polychaetes, Osinga *et al*., 2001; Ribeiro *et al*., 2003; Rützler, 2012), and microorganisms (including viruses, bacteria, archaea, microalgae, and other microeukaryotes, Thomas *et al*., 2016; Pita *et al*., 2018). Sponges share intricate interactions with their microbiome, acting together as a functional unit known as the holobiont (Webster and Thomas, 2016). The host-microbiome interactions ensure diverse ecological processes for the holobiont, such as organic matter cycling, protection against oxidative stress, or production of chemical defenses and vitamins (Regoli *et al*., 2000; Webster and Thomas, 2016; Pita *et al*., 2018; Han *et al*., 2019).These diverse interactions contribute to sponge resilience under future ocean conditions, including seawater warming (Bennett *et al*., 2017; Posadas *et al*., 2022) and ocean acidification (Ribes *et al*., 2016). Additionally, marine sponge microbiomes provide stability and essential adaptations for their host facing extreme trophic conditions, allowing survival in nutrient-poor environments (Rix *et al*., 2017), and under eutrophic pressures (Simister *et al*., 2012; Baquiran and Conaco, 2018). However, this contribution varies across different regions and sponge species, requiring further investigation in multiple habitats and understudied species (De Castro-Fernández *et al*., 2023). In particular, the role and dynamics of sponge microbiomes in freshwater ecosystems remain poorly understood, especially given the high level of anthropogenic impact in these habitats (Lo Giudice and Rizzo, 2024).

Freshwater sponges (Order Spongillida) comprise approximately 270 species (Pronzato *et al*., 2017, de Voogd et al., 2025) and are widely distributed across freshwater bodies worldwide (except Antarctica) occurring in diverse habitats such as rivers, lakes, ponds, and urban canals (Manconi and Pronzato, 2008). Their broad distribution results from adaptations to a variety of environmental conditions, including a wide range of temperatures, light levels, pH, oxic conditions, and pollutant concentrations (Manconi and Pronzato, 2008). In light-exposed habitats, freshwater sponges generally exhibit a green coloration resulting from colonization by microalgal photosymbionts. These microalgae, such as *Chlorella* spp., can provide a significant source of carbon and energy for the host, contributing up to 20% of photosynthate production (Wilkinson, 1980). The symbiosis between *Chlorella*-like microalgae and their sponge hosts, also involves key recognition mechanisms by the host immune system, through cellular and genetic responses, as observed in the freshwater sponge *Ephydatia muelleri* (Hall *et al*., 2021). However, this association between freshwater sponges and their photosymbionts can be disrupted, leading to a dysbiotic state, as observed with the brown rot syndrome of the Baikal sponge *Lubomirskia baikalensis* (Kulakova *et al*., 2018; Belikov *et al*., 2019). Significant changes in the bacterial communities have been observed in these diseased sponges, with the potential emergence of bacterial pathogens. Nevertheless, the role of these bacterial community shifts and associated environmental parameters in relation to brown rot syndrome remains complex and not yet fully understood (Kulakova *et al*., 2018). Recently, two studies have investigated the specific effects of environmental parameters on bacterial communities associated with several freshwater sponge species (Keleher *et al*., 2025; Rizzo *et al*., 2025). The first study, conducted on *E. muelleri* and *Spongilla lacustris* in the sub-Artic Psvik river, revealed high concentrations of bioaccumulated chemical contaminants (particularly chemicals of emerging concern) within sponge tissues, which may explain the observed spatial differences in bacterial communities (Rizzo *et al*., 2025). The second study, conducted on *Radiospongilla crateriformis*, *Trochospongilla* sp. and *Eunapius fragilis* in North Carolina also identified spatial patterns in microbiome composition, in addition to temporal differences relative to the surrounding freshwater (Keleher *et al*., 2025). Together, these two studies indicate that the health and physiology of the freshwater sponge hosts are essential factors to consider, including stress related to chemical contaminants (Rizzo *et al*., 2025), as well as different life stages during the growing season and period of tissue degradation (Keleher *et al*., 2025).

*Spongilla lacustris* (Family Spongillidae) is a freshwater sponge species widely distributed in temperate regions, especially in Europe (Manconi and Pronzato, 2008) and hosts *Chlorella*-like photosymbionts (Jensen and Pedersen, 1994). Such symbionts can be lost after the sponge growth, particularly during the cold period, through degradation or expulsion of algal cells, resulting in “bleached” sponge tissues ranging from white to brownish (Williamson, 1979). In addition to these photosymbionts, a large bacterial diversity can be observed within *S. lacustris*, dominated by several phyla including Alphaproteobacteria, Bacteroidota, Actinobacteria, Betaproteobacteria and Gammaproteobacteria (Gernert *et al*., 2005; Graffius *et al*., 2023; Paix *et al*., 2024). Before the cold period in winter, *S. lacustris* produce dormant cysts, known as gemmules for asexual reproduction (Gilbert, 1975). During this process, termed gemmulation, adult sponges transmit bacterial communities both to the surface of the gemmule coating and within the gemmules alongside the dormant cells (Paix *et al*., 2024). Interestingly, this horizontal transmission of bacterial symbionts appears to be essential for holobiont stability during the early growth stages of juveniles. Together with Keleher *et al*., (2025), these studies highlight the importance of considering the freshwater sponge development periods to better understand temporal dynamics within the holobiont. Current knowledge of microbial dynamics during the adult phases of *S. lacustris*, such as gemmulation, bleaching and senescence, is still limited.

In the present study, a large population of *S. lacustris* was observed around the canals of Leiden (the Netherlands) from October to December 2021, providing an ideal context to investigate its associated bacterial community under contrasting environmental conditions and across adult life stages. This study aims to examine how the bacterial composition and diversity associated with *S. lacustris* vary across space and time, and to identify the major ecological factors driving these dynamics. We hypothesize that bacterial communities are strongly linked to pronounced changes in the physiological state of *S. lacustris* during the mature phase of its life cycle, as well as freshwater physico-chemical conditions, particularly in urbanized environments. Accordingly, this study employed spatio-temporal *in situ* monitoring across multiple conditions to simultaneously account for distinct physiological stages of the adult sponge cycle (gemmulation, bleaching and senescence), and the influence of contrasting environmental parameters, including temperature, pH, turbidity, and nutrient concentrations (phosphates and ammonium).

## Material and methods

### Sampling strategy, biological materials, and measurement of environmental parameters

Fieldwork was conducted at six sampling times (T1 to T6) at two-week intervals between October 2021 and December 2021 at three sites in South-Holland province (The Netherlands, **Table S1**). These sites were selected based on their contrasting levels of anthropization (**Figure 1A**). Specifically, site S1 (52.154282 N, 4.484189 E) was located in a canal within the city of Leiden, where the anthropogenic influence is expected to be the highest. Site S2 (52.191463 N, 4.498750 E) is located in a small harbor of the village Warmond, which is connected to the recreational lake Kaag. Site S3 was located in a canal of the peri-urban area of the village of Oegstgeest, directly connected to the Rhine River (52.10241N, 4.27017E).

**Figure 1.**
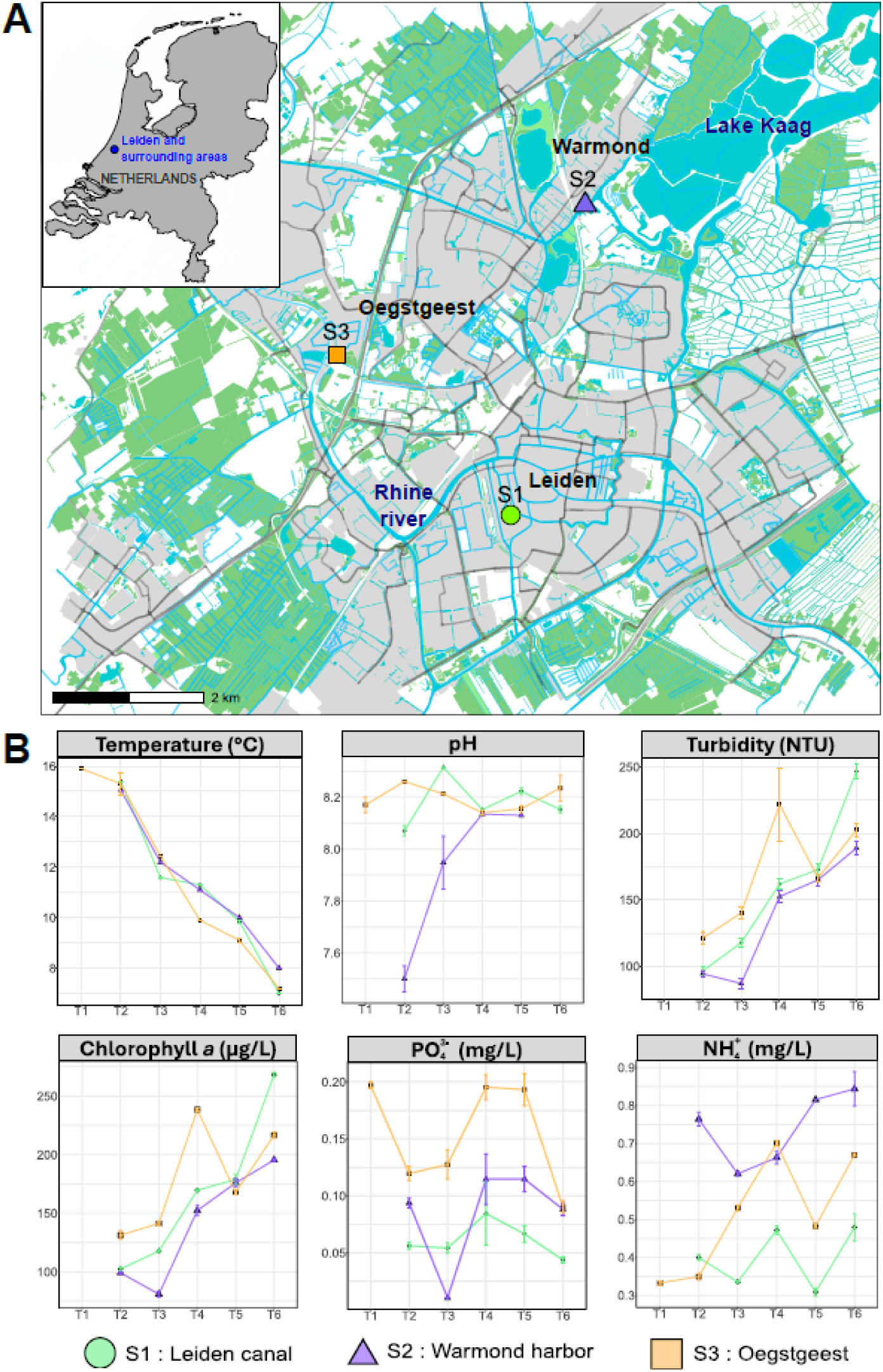
Sampling sites of the study. (A) Maps of the sampling site locations. Farmlands and grasslands are represented in green, residential areas and highways are represented in grey, while canals, natural waters and reservoirs are represented in blue. (B) Temporal variations of temperature, pH, turbidity, chlorophyll *a* concentration, and nutrient concentrations (PO_4_^3-^ and NH_4_^+^), within each site.

At each sampling time, four replicate sponges were collected at depths between 0 and 1 m. Samples from T1 were collected only at site S3, resulting in a total of 63 samples (n=3 for site S2 at T6, due to the limited number of specimens observed, **Table S1**). In addition, two to four replicates of surrounding freshwater (400 mL) were collected for metabarcoding analyses, yielding 60 additional samples (**Table S1**). After collection, sponges were cut using a sterilized scalpel blade and stored at −20°C in 5 mL of absolute ethanol (EtOH) until metabarcoding analyses. Freshwater samples for metabarcoding were filtered through 0.2 µm pore size filters (FP 30/0.2 CA-S, Whatman GMbH) and stored in 600 µL of hexadecylcetyltrimethylammonium bromide (CTAB) buffer at −20°C until processing. Additional freshwater samples (30 mL) were collected for nutrient analyses, filtered through 0.2 µm membrane filters (FP 30/0.2 CA-S, Whatman GMbH), and frozen at −20°C. For morphological analyses, sponge tissues were photographed prior to cutting and stored at room temperature in 96% EtOH.

Temperature and pH were measured *in situ* using a multiparameter Hanna HI98494 probe (Hanna Instruments). Turbidity and chlorophyll *a* concentrations were measured *in situ* using an AquaFluor® (Turner Designs) for all sampling except T1. Nutrient concentrations (NH_4_^+^ and PO_4_^3-^) in freshwater samples were quantified using Spectroquant® ammonium and phosphate test kits following the manufacturer’s instructions (Merck). Measurements were performed using a Spark® Multimode Microplate Reader (Tecan).

### Identification of *S. lacustris* and morphological description

The identification of *S. lacustris* specimens was performed based on examination of skeletal elements and gemmoscleres. A voucher specimen was deposited in the sponge collection of Naturalis Biodiversity Center (RMNH.POR.12472).

The gemmulation life stage of the sponges was identified by estimating the relative mass of gemmules within the whole sponge tissue. For each sample, three pieces of sponge were subsampled (i.e., one encrusting part, and two from the middle and apical parts of the branches), and gemmules were separated from sponge tissues following the protocol described by Leys *et al*., (2019). After drying, gemmules were weighed, and their relative mass was calculated as a percentage of the total sponge mass. The presence of bleached and diseased tissues was also analyzed based on coloration, ranging from green color (indicative of photosymbiont presence) to brownish in bleached specimens, and blackened in degraded tissues (**Figure S1).**

### DNA extraction, library preparation, and High Throughput Sequencing (HTS) of the 16S rRNA gene amplicons

Approximately 0.1-0.2 g of each sponge sample was cut for DNA extraction, using sterilized tweezers and scalpel blades. The FastDNA™ SPIN Kit for Soil (MP Biomedicals, Inc.) was used according to the manufacturer’s instructions. For freshwater samples, the CTAB DNA extraction protocol for PCTE filters was used according to Turner *et al*., (2014). One extraction blank was prepared for each sample type (sponge and freshwater samples).

The library preparation for metabarcoding analyses was conducted through a two-step PCR protocol for all samples, including sponge and freshwater samples, extraction blanks and PCR1 negative controls. For PCR1, the V3-V4 region of the 16S rRNA gene was targeted with the 341F (CCTACGGNGGCWGCAG) and 785R (GACTACHVGGGTATCTAATCC) primers (Klindworth *et al*., 2013). PCR1 reactions were performed using the KAPA HiFi HotStart Ready Mix PCR Kit (Roche Molecular Systems, Inc.), in a T100 Thermal Cycler (Bio-Rad, Hercules, CA, United States). The following thermal cycling scheme was set up: initial denaturation step at 95°C for 3 min, followed by 30 cycles of denaturation step at 98°C for 20s, annealing at 57°C for 30 s, and extension at 72°C for 30 s, followed by the final extension step at 72°C for 1 min. PCR1 products were checked on E-GEL™ (agarose gels at 2%, Invitrogen), and the absence of amplification was validated for the negative controls and the two extraction blanks. PCR1 products were then cleaned using NucleoMag NGS-Beads (bead volume at 0.9 times the total volume of the sample, Macherey Nagel, Düren, Germany) and the VP 407AM-N 96 Pin Magnetic Bead Extractor stamp (V&P Scientific, San Diego, CA, United States).

PCR2 were performed using IDT xGen™ NGS Adapters & Indexing Primers kit (Integrated DNA Technologies, Inc.) with an initial denaturation step of 3 minutes at 95°C followed by 8 cycles of 20 seconds at 98°C, 30 seconds at 55°C and 30 seconds at 72°C, and a final extension step of 5 minutes at 72°C. Successful labeling of PCR2 products was then checked with the Fragment Analyzer Agilent 5300 using the DNF-910-33 dsDNA Reagent Kit (35–1,500 bp) protocol (Agilent Technologies, Santa Clara, CA, United States) and concentration was determined with PROSize 3.0 software. Using the QIAgility (Qiagen, Hilden, Germany), samples were pooled together at equimolar concentration. The pool was then cleaned using NucleoMag NGSBeads and the DNA concentration was quantified using Tapestation 4150 (Kit HSD 5000, Agilent Technologies, Santa Clara, CA, United States). The amplicon pool was sent to BaseClear (BaseClear B.V., Leiden, The Netherlands) for MiSeq Illumina sequencing (V3 2*300 PE platform).

### 16S rRNA gene metabarcoding data processing and analyses

The raw reads of all samples were initially processed by BaseClear B.V. for demultiplexing (using 274 bcl2fastq version 2.20, Illumina), and filtering based on two quality controls (using Illumina 275 Chastity filtering, and a PhiX control signal filtering). Amplicon sequences were then processed following the DADA2 pipeline under RStudio (dada2 R package, https://benjjneb.github.io/dada2/tutorial.html), allowing an inference to Amplicon Sequence Variant (ASV) (Callahan *et al*., 2016, 2017). Sequences were filtered and trimmed for forward and reverse reads at truncation length of 280 and 230 base pairs, respectively, maxN = 0, maxEE = 2, and truncQ = 2. After constructing the ASV table, chimeric sequences were filtered out and the taxonomic assignment from domain to genus levels was performed using the Silva v138 reference database (Quast *et al*., 2013). The ASV and taxonomy tables produced by the pipeline were then combined into a phyloseq object, together with the sample metadata table, using the “phyloseq” R package (McMurdie and Holmes, 2013). 671 ASVs assigned as Eukaryotes, chloroplasts, or mitochondria were removed representing 3.2% of the total ASVs. The dataset was then cleaned from contaminants with the negative controls and extraction blanks using the “decontam” R package, with the “prevalence” method (Davis *et al*., 2018) resulting in 19 ASVs removed. For subsequent analyses, one sponge replicate (at S3T5) and four freshwater samples (at S1T2, S1T3, S2T3, and S3T3) were removed from the analysis (**Table S1**), due to their limited sequencing depth (< 7000 reads).

Rarefaction curves are plotted through the “vegan” R package (Oksanen *et al*., 2019). The α-diversity metrics were estimated using Chao1 (estimated richness), Pielou (evenness), and Shannon (both richness and evenness) indices on the rarefied dataset (rarefaction performed to the minimum library size, i.e. 16388 reads for the sponge samples and 8165 for the freshwater samples) with using “vegan” and “phyloseq” R packages. Differences in α-diversity measures across the different sample types, sites, and times were tested according to the normality of their distribution (assessed with a Shapiro test), with ANOVA test followed by HSD Tukey’s post-hoc test, or a Kruskal-Wallis test followed by a pairwise Wilcoxon test. Following recommendations for β-diversity and compositional analyses from McMurdie and Holmes, (2014); Gloor *et al*., (2017), all other analyses were conducted without rarefaction, using the datasets normalized to the total number of sequences per sample. β-diversity was analyzed with a Non-metric Multidimensional Scaling (NMDS), using the Bray-Curtis dissimilarity. NMDS analysis was also plotted using branch colors together with relative gemmule weight fitted as a continuous variable using the *ordisurf* function from the “vegan” package. Differences in β-diversity between groups were statistically checked with one-way PERMANOVA tests, followed by pairwise adonis tests from the “pairwiseAdonis” R package (Martinez Arbizu, 2020). β-diversity dispersion analysis was performed using the *betadisper* function from the “vegan” package. To analyze the environmental parameters (temperature, turbidity, concentration of chlorophyll *a*, pH, concentration of phosphates and ammonium) as explanatory factors on the β-diversity, a db-RDA was performed, and the significance of the model was tested using the function *anova.cca*() from the “vegan” R package. As chlorophyll *a* and turbidity were found to be highly redundant, only chlorophyll *a* measures were kept for the analysis. The relative contribution of the environmental parameters as explanatory factors of the β-diversity variance was calculated with the variance partitioning using the *varpart* function from the “vegan” package with the formula [∼Turbidity + Chlorophyll *a*, ∼Temperature, ∼ pH + NH_4_^+^ + PO_4_^3-^)].

The sponge core community was analyzed with the *core* function from the “microbiome” R package (Lahti *et al*., 2017). The core community was defined with a prevalence threshold of 90% for all sponge samples (representing here at least 56 out of the 62 sponge samples), as suggested in Maslin *et al*., (2024). The relative richness of core ASVs within each sample was estimated by comparison with the total richness.

For differential analyses, phylogenetic heat trees were plotted using the “metacoder” R package (Foster *et al*., 2017), to identify the taxa differentially abundant within each group, according (i) to the sponge life stage, and (ii) sampling sites. For a better readability of the graphical output, the differential analyses were performed with a dataset excluding rare ASVs (relative abundance < 0.04%).

## Results

### Environmental conditions and morphological description of *S. lacustris* samples

Water temperatures decreased from 16°C to 7°C from October to December (**Figure 1B)**, without significant differences between sites (ANOVA test: *p* > 0.05). Chlorophyll *a* and turbidity both increased significantly with time (ANOVA test: *p* < 0.001), without any significant differences between sites (ANOVA test: *p* > 0.05). Significant differences in pH were observed between sites (Kruskal-Wallis test: *p* < 0.05), with lower values for S2, especially at T2 and T3. Significant differences between sampling sites were also observed for ammonium (NH_4_^+^) concentrations (ANOVA test: *p* < 0.001), with higher values for S2 and lower concentrations for S1, and phosphate (PO_4_^3-^) concentrations with higher values for S3 (ANOVA test: *p* < 0.001).

Different morphological characteristics of the sponge samples were studied with observations of the presence of gemmules, bleached tissues, and blackened tissues. The bleaching of the sponge tissues was mainly observed after T3, regardless of the site (**Figure S1A, Table S1**). From T4 to T5, most of the sponge specimens were partially bleached, with the presence of photosymbionts occurring only within the light-exposed side of the branches. Consequently, sponge samples were divided into three different groups: (i) green sponges, (ii) partially bleached, and (iii) fully bleached. Additionally, several sponge specimens were observed with blackened tissues after T3 (especially for T6 samples), regardless of the sampling sites (**Figure S1B)**. Almost no gemmules were observed within sponge tissues from T2 to T3, while the relative weight of the gemmules within the tissues increased from T4 in S1 and T5 for S2 and S3 (**Figure S2**). The highest relative weights were observed in T6 for each site (between 11% and 34% for S1, 10% to 25% for S2, and 9.5% to 22%).

### α- and β-diversity analyses of planktonic and sponge-associated bacterial communities

After filtration of the reads, rarefaction curves reached a plateau for all samples (**Figure S3**). Chao1, Pielou, and Shannon indices revealed higher values for the planktonic communities compared to sponge-associated ones (**Figure S4**). When focusing on sponge samples (**Tables S2 and S3**), significant differences between sites were observed, with lower values for S1 compared to S2 for Chao1 (**Figure S4, Table S3**), and lower values for S1 compared to S3 for Shannon (**Figure S4, Table S3**). No significant differences between the sites were observed for the Pielou index. Within S1 and S2, significant differences in Shannon index were observed over time, with significantly lower values at T6 compared to T3 for S1, and lower values at T5 compared to T3 and T4 for S2 (**Figure S4**).

The NMDS plot and PERMANOVA test performed with all samples revealed a significant difference between planktonic and sponge-associated bacterial communities (**Figure 2A**, **Table S4**). When focusing specifically on sponge samples, the NMDS plot and PERMANOVA test revealed a significant temporal shift on the first NMDS axis (**Figure 2B**, **Table S5**). The multivariate pairwise adonis test confirmed these temporal differences observed between each sampling time, except between T1 and T2 (**Table S6**). Spatial differences can also be observed on the second axis of the NMDS plot (NMDS2), with S2 and S3 samples clustered separately (**Figure 2B**). These spatial differences were confirmed by the PERMANOVA test (**Table S5**), however, adonis pairwise comparisons between each site were not significant (**Table S7**). The NMDS analysis was also visualized according to sponge morphology, using branch coloration, with relative gemmule weight fitted as a continuous variable (**Figure 2C**). A distinct cluster of green sponges lacking gemmules, corresponding to samples from T1, T2, and T3 were observed. The overall shift in β-diversity along the temporal axis (NMDS1 axis) was consistent with increasing gemmule weight and progressing sponge bleaching (**Figure 2C**). Additionally, the NMDS analysis revealed higher heterogeneity in β-diversity distances among fully bleached sponges with high relative gemmule weight, primarily associated with samples from T5 and T6, compared to green samples from T1 to T3 (**Figure 2B** and **2C**). This pattern was confirmed by β-dispersion analysis, which showed significant differences between sample times (**Table S8**), with a general increase in distance to centroids from T4 to T6 across all sites (**Figure 2D**). No significant differences in distance to centroids were detected between sites (Kruskal-Wallis tests: *p* > 0.05). Freshwater samples exhibited lower β-dispersion values compared to sponge samples (T-test: *p* <0.001), with a general temporal trend displaying a U shape, for which the highest values occurred at T1, T2 and T6 across all sites (**Figure 2D**).

**Figure 2.**
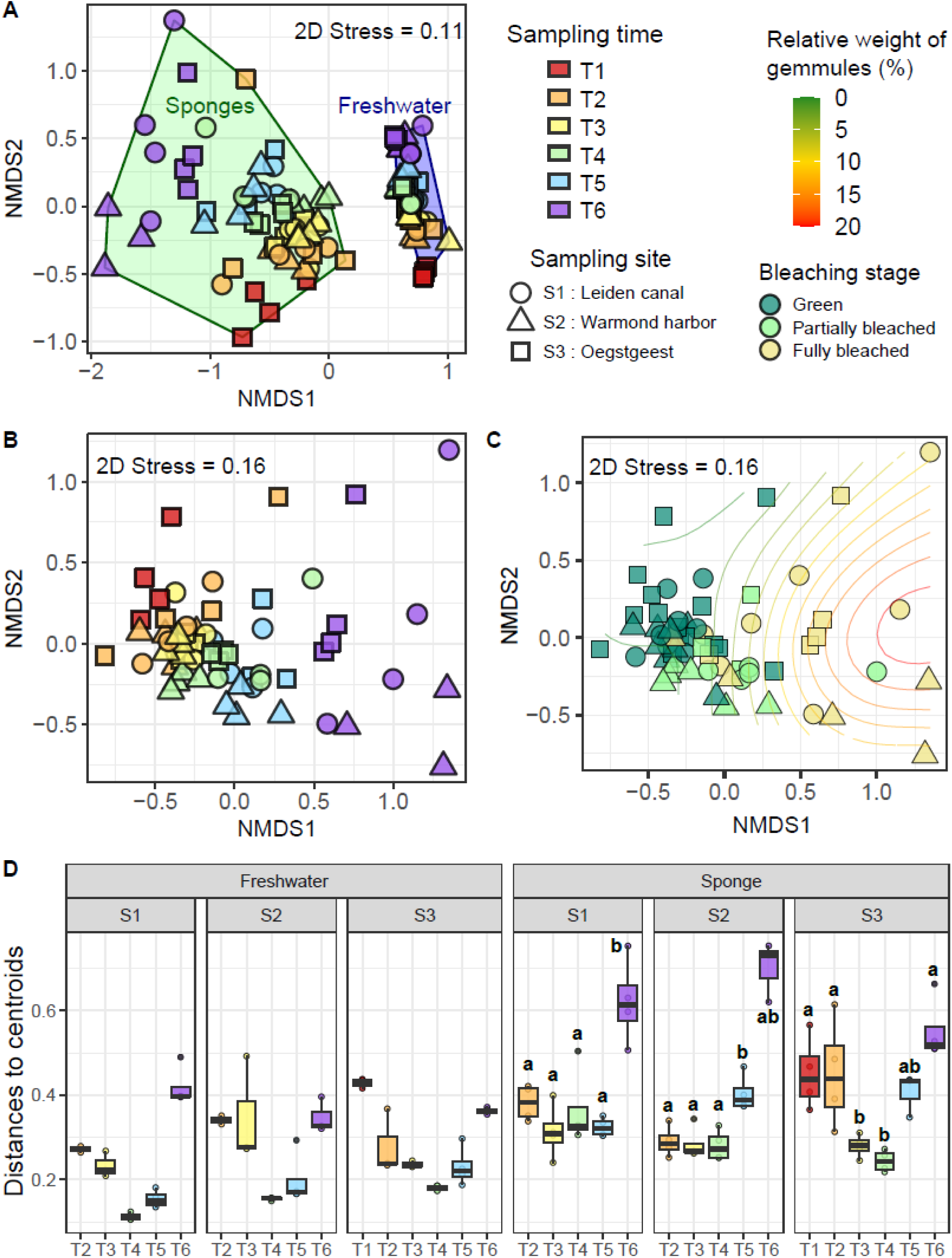
β-diversity analyses of the freshwater and sponge-associated bacterial communities. **A.** NMDS plot of freshwater and sponge samples. **B.** NMDS plot of sponge samples displaying the sample sites and sampling time. **C. N**MDS plot of the sponge samples displaying the sponge colors and the relative weight of the gemmules. Curved lines represent the fitting of the relative weight of the gemmules using the ordisurf function. **D.** Dispersion of the β-diversity of the bacterial community of freshwater and sponge samples for each site and time.

The distance-based redundancy analysis (db-RDA) was performed using the Bray–Curtis dissimilarity index to assess environmental parameters as explanatory variables of β-diversity (**Figure 3A**, ANOVA.CCA: *p* < 0.001). Opposing temporal trends in temperature and chlorophyll *a* concentration (the latter being redundant with turbidity) were the main drivers of temporal β-diversity dynamics particularly across samples from T2 to T6. Additionally, higher ammonium concentrations, pH, and to a lesser extent, phosphate concentrations contributed to spatial differences in β-diversity, with a cluster specifically associated with samples from site S2. Variance partitioning analysis indicated that 5% of the β-diversity variance was explained by turbidity and chlorophyll *a* concentration, 1% by temperature, and 5% by nutrient concentrations (NH_4_^+^ and PO_4_^3-^) together with pH (**Figure 3B**). When combined, turbidity, chlorophyll *a* concentration, and temperature accounted for 9% of the variance. Combined, all environmental parameters explained 3% of the total β-diversity variance (**Figure 3B**).

**Figure 3.**
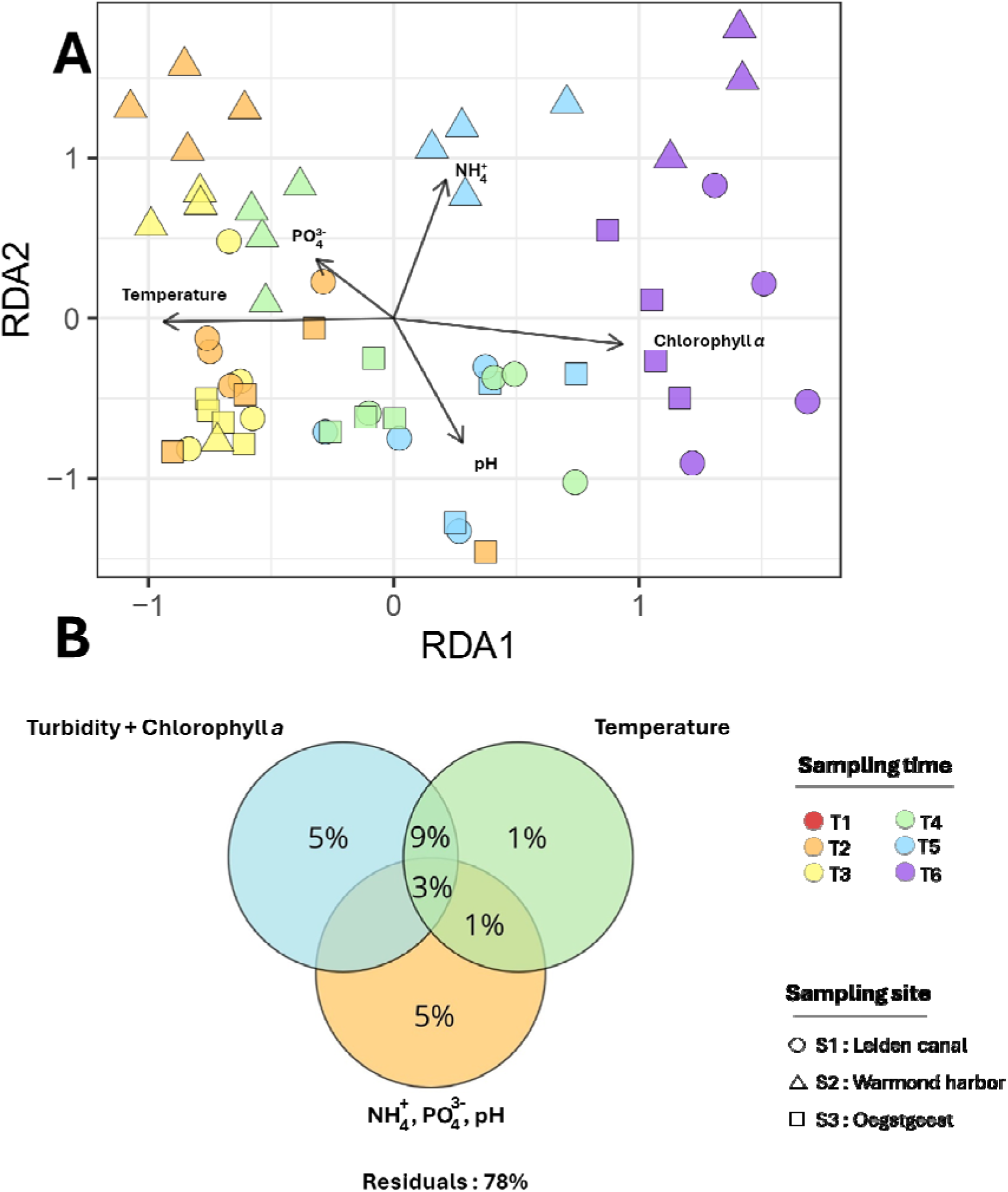
Analyses of the environmental variables as explanatory factors of the bacterial community β-diversity within sponge samples. **A.** db-RDA score plot. **B.** Venn diagram showing the variance partitioning.

### Composition of the overall and core bacterial community

Although two dominant families were shared between sponge and freshwater samples (*Comamonadaceae* and *Sporichthyaceae*), pronounced differences in bacterial community composition were observed between these two sample types (**Figure S5**). Alphaproteobacterial taxa were more diverse and abundant in sponge samples, with families such as *Elsteraceae*, *Terasakiellaceae*, *Rhodobacteraceae*, and *Sphingomonadaceae* being absent from planktonic communities. Gammaproteobacterial families were also more diverse in sponges, including *Methylomonadaceae*, *Pseudomonadaceae*, *Spongiibacteraceae*, *Rhodobacteraceae,* and *Xanthomonadaceae*, which were not detected in planktonic communities. In contrast, *Paenibacillaceae* (Firmicutes), along with several families belonging to the Chloroflexi and Campylobacterota phyla, were specifically observed in planktonic communities **(Figure S5**).

Across all conditions, sponge-associated bacterial communities were dominated by several families, including the *Chitinophagaceae* (15.68%; SD = ±0.07), the *Elsteraceae* (8.5%; SD = ±0.06), the *Comamonadaceae* (10.37%; SD = ±0.04), and the *Terasakiellaceae* family (8.5%; SD = ±0.06) (**Figure S5A**). Across the three sites, a general decrease in families belonging to the phylum Actinobacteriota, as well as in *Terasakiellaceae* was observed over time, whereas Bacteroidota families increased and dominated the community at T6.

Core community analysis of *S. lacustris* identified a total of 20 core ASVs. The relative abundance of these ASVs ranged from 12.93% to 82.04% (average: 55.55%, SD: ±13.13) (**Figure 5**). The dominant core families belonged to *Chitinophagaceae* (Bacteroidota phylum), *Elsteraceae*, and *Terasakiellaceae* (Proteobacteria phylum). No significant differences in core ASV relative abundance were observed among sampling sites or sampling times (ANOVA test: *p* > 0.05). The relative richness of core ASVs per sample ranged from 2.26% to 16.83% (average: 8.81%, SD = ±2.67, **Figure S6**). Significant differences were detected among sites (**Table S9**), with lower values at site S2 compared to sites S1 and S3 (**Figure S6**). No significant temporal differences were observed, except at site S2, where relative richness was lower at T6 than at T5.

### Differential analyses of the sponge associated bacterial communities

Differential analyses based on bleaching stages of the sponges first revealed a high diversity of bacterial taxa that were either specific to green or to bleached sponges (**Figure 4**). More specifically, taxa belonging to the phylum Actinobacteriota were found at higher relative abundances in green sponges, particularly within families *Sporichthyaceae, Ilumatobacteraceae*, and *Microbacteriaceae*. Similar patterns were observed for several other phyla including Bacteroidota (e.g. *Solitalea*, *Fluviicola*, *Rurimicrobium,* and *Parasediminibacterium*), Proteobacteria (*Pseudohongiellaceae* and *Methylophilaceae* families, as well as the genera *Malikia*, *Alsobacter* and *Sphingorhabdus*), Nitrospirota (*Nitrospira*), Campylobacterota (*Pseudarcobacter*), Armatimonadota (*Armatimonas*), Spirochaeota (*Leptospiraceae*) and Verrucomicrobiota (*Luteolibacter*). In contrast, fully bleached specimens were specifically associated with the genera *Defluviimonas*, *Tabrizicola* (*Rhodobacteraceae* family), Nitrospira (*Nitrospiraceae* family), *Neisseriaceae* (Burkholderiales order), and *Flavobacterium* (*Flavobacteriaceae* family).

**Figure 4.**
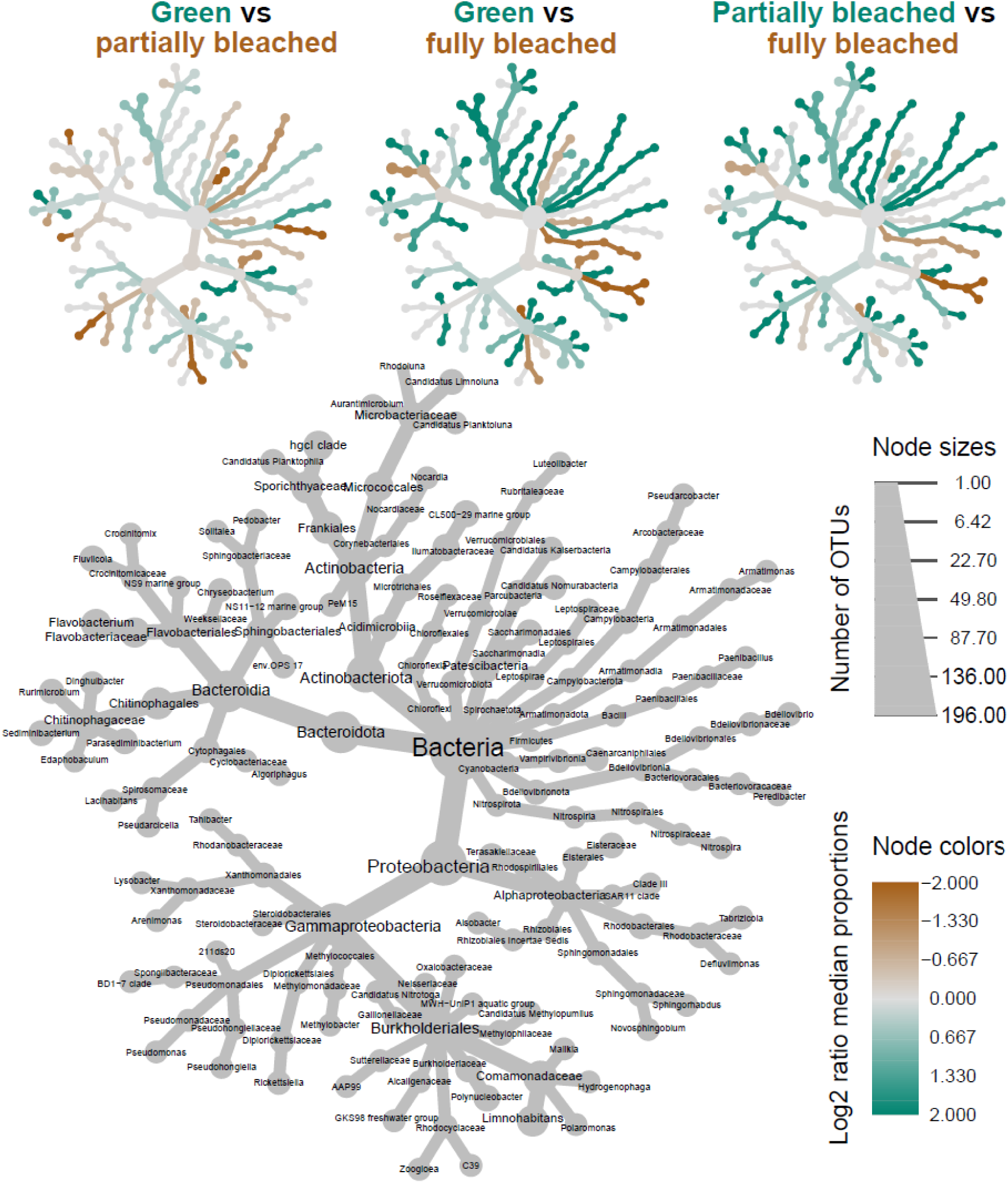
Phylogenetic heat trees representing the bacterial taxa significantly and differentially abundant between the green, partially bleached, and fully bleached sponges. For each taxon, (i) the colors of their associated nodes correspond to the log2 fold change between the sponge coloration groups, (ii) the size of the nodes corresponds to the relative abundance of each taxon.

Differential analyses were also conducted among sites to identify taxa with significantly different relative abundances specific to (i) the ammonium-rich site (S2), and (ii) the phosphate-rich site (S3). The first analysis revealed significantly higher relative abundances of Saccharimonadales and *Methylobacter* (family *Methylomonadaceae*, class Gammaproteobacteria) at site S2 compared to sites S1 and S3 (**Figure S7A**). In contrast, sites S1 and S3 showed higher relative abundance of members of the *Leptospiraceae* family (**Figure S7A**). Additionally, the genera *Rurimicrobium* (family *Chitinophagaceae*) and *Malikia* (family *Comamonadaceae*) were significantly enriched at site S3, whereas sites S1 and S2 were enriched in taxa belonging to the order Saccharimonadales (**Figure S7B**).

**Figure 5.**
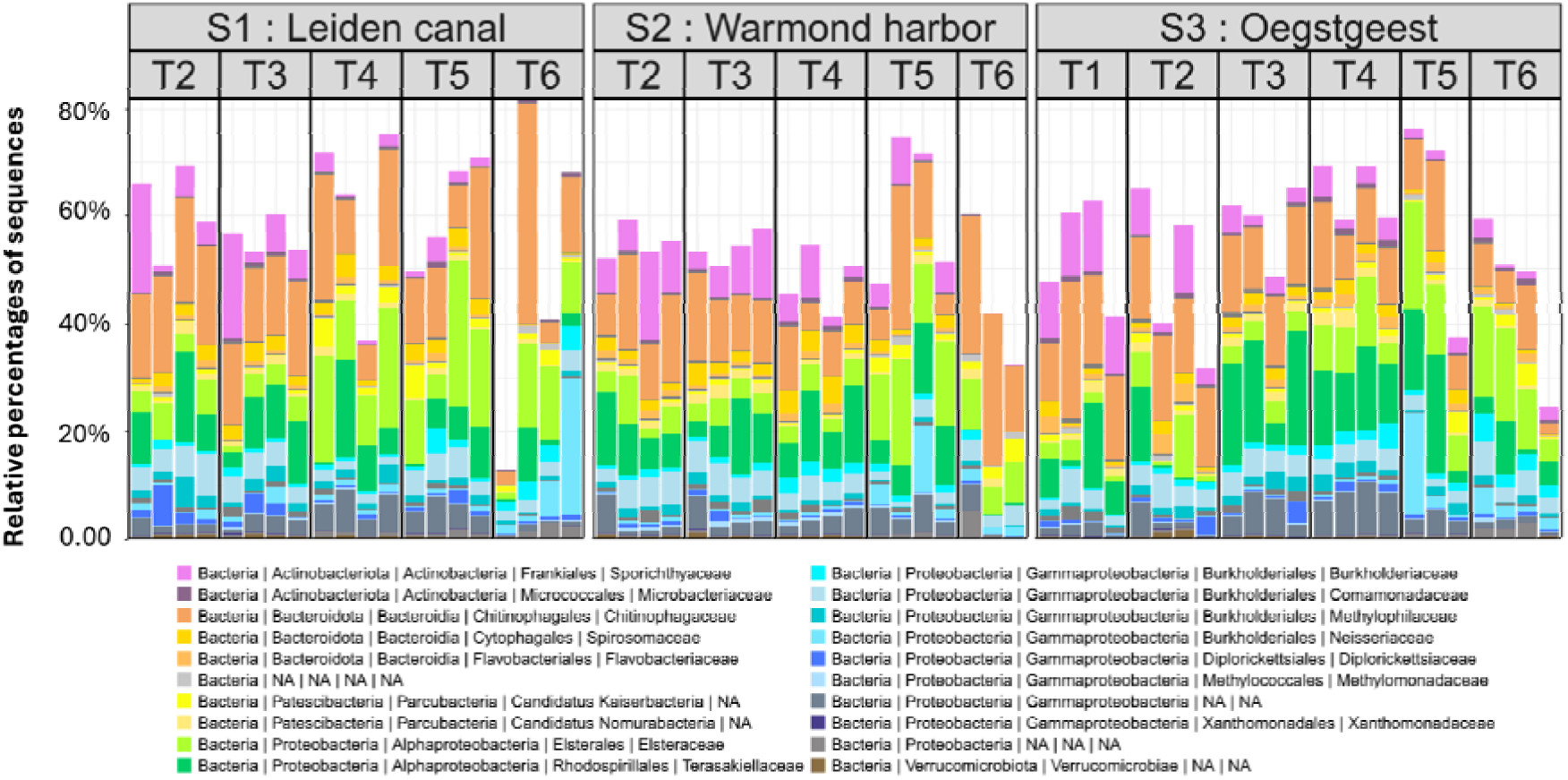
Barplots of the relative abundance of the bacterial core community composition (at the family level) in *S. lacustris* occurring at least 90% of the samples. “Other” and “NA” correspond to families with a relative percentage below 5% and unaffiliated families, respectively.

## Discussion

This study aimed to identify the main factors driving the spatiotemporal dynamics of bacterial community diversity associated with the freshwater sponge *Spongilla lacustris*. We hypothesized that both anthropogenic pressures and host life cycle-related physiological factors shape the bacterial microbiome of *S. lacustris*. In addition, changes in bacterial community composition were considered as a potential factor influencing host health over time. To evaluate these effects, sponge samples were collected from three sites characterized by contrasting levels of anthropogenic pressures. Sampling was conducted during a period when the physiology of the adult specimens underwent marked transitions, from healthy, green-colored sponges to fully bleached and degraded individuals.

### Spatial differences in the bacterial diversity of *S. lacustris*, explained by contrasted nutrient concentrations

The first step of this study was to validate the selection of sampling sites by confirming that they reflected contrasting levels of anthropogenic pressures, based on measurements of chemical parameters such as nutrients concentrations. Site 1, located within the canals of Leiden city center, was initially considered to be the most anthropized site and was expected to exhibit the highest nutrient concentrations. In contrast, sites S2 and S3 were considered to have greater hydrological connectivity. Specifically, site S2 was located in Warmond Harbor, and connected to Lake Kaag, while site S3 was situated in a peri-urban area and directly connected to the river Rhine. Unexpectedly, the highest ammonium concentrations were measured at site S2, whereas the highest phosphate concentrations were observed at site S3. The elevated phosphate levels at site S3 may be attributed to its downstream position relative to Leiden and its canals network, potentially promoting nutrient accumulation. In addition, local anthropogenic pressures near site S3, such as the presence of a major highway and an industrial zone may further contribute to phosphate inputs. Site S2, located in Warmond Harbor and connected to the recreational lake Kaag, is also situated near the Leiden Noord wastewater treatment plant (52.1744282 N, 4.4944329 E), which may discharge nutrient-rich effluents. Together, these anthropogenic influences may promote eutrophication, explaining the elevated ammonium concentrations observed at these sites.

Our study highlights the importance of nutrient concentrations as key environmental drivers underlying spatial differences in bacterial β-diversity associated with *S. lacustris*. The respective effects of ammonium and phosphate could be distinguished, as distinct bacterial communities were observed at sites S2 and S3. Site S2 was characterized by elevated ammonium concentrations and a specific enrichment of Saccharimonadales*, Pseudoarcobacter* and *Methylobacter*. Saccharimonadales and *Pseudoarcobacter.* There taxa have been associated with nitrifying or denitrifying assemblages and typically proliferate in sewage, wastewater effluents, and brackish environments with high nitrogen loads (Carey and Migliaccio, 2009; Aoki *et al*., 2023; Wang *et al*., 2023; Qian *et al*., 2024; Emmanuel *et al*., 2025). Their presence within *S. lacustris* at site S2 is therefore consistent with the strongly eutrophic conditions and elevated ammonium concentration observed at this site. These taxa may reflect the key role of ammonium in the nitrogen cycling of the *S. lacustri*s holobiont, as previously reported for marine sponge holobionts in Caribbean reefs (Gantt *et al*., 2019). The enrichment of *Methylobacter* further supports the influence of urban eutrophication at site S2. This type I methanotroph, commonly found in freshwater lakes (Borrel *et al*., 2011), has also been linked to methanotrophic biofilms in Dutch urban canals (Pelsma *et al*., 2023), where methane emissions can co-occur with diffuse nutrient pollution. In contrast, higher phosphate concentrations were measured at site S3, where *Malikia* and *Rurimicrobium* were identified as biomarkers genera. *Malikia* has been proposed as an indicator of wastewater pollution, partly due to its denitrification capacity in nutrient-enriched waters (Ruprecht *et al*., 2021). *Rurimicrobium*, originally isolated from nutrient-rich agricultural soils (Dahal *et al*., 2017) similarly suggest adaptation to nutrient-rich environments, as also observed in aquaponic systems (Kasozi *et al*., 2020).

The distinct enrichment of these biomarker taxa at both sites S2 and S3 suggests that these bacterial groups may respond positively to eutrophication-related conditions specific to each site. While ammonium and phosphate clearly emerged as important drivers of β-diversity in *S. lacustris*, focusing solely on these two nutrients likely underestimate the complexity of environmental factors shaping *S. lacustris* microbiomes. Because 78% of the variance remained unexplained by the measured environmental parameters, a broader range of chemical drivers should be considered in future studies. For instance, relationships between freshwater sponge microbiome and dissolved or particulate organic carbon (DOC, POC) could be examined, as has been done for Caribbean coral reef sponges, where no clear associations were detected (Gantt *et al*., 2019). In addition, nitrogen species beyond ammonium, including nitrate and nitrite, should be considered in light of key nitrogen cycling pathways mediated by sponge-associated microbiomes, such as nitrification and denitrification, as reported for deep-sea sponge holobionts (Radax *et al*., 2012; Tian *et al*., 2016; Glasl *et al*., 2024). Other important water quality drivers include pharmaceutical compounds classified as chemicals of emerging concern, which has been shown to bioaccumulate in freshwater sponges such as *S. lacustris* and *E. muelleri* (Rizzo *et al*., 2025). Integrating these additional parameters into future studies, together with functional approaches such as metagenomics, metatranscriptomics or metametabolomics, would provide deeper insights into how multiple stressors influence microbial activities within *S. lacustris* across spatially heterogeneous environments. As an initial step, targeted investigation of DOC assimilation and methane oxidation pathways potentially linked to *Methylobacter* at site S2 would be particularly informative.

However, even if differences of bacterial β-diversity appeared on the three sites with differentially abundant taxa specific to S2 and S3, no clear spatial patterns were noticed for the sponge morphology, α-diversity and the core community composition. Those observations on the microbiome might be explained by a prevailing influence of the observed temporal parameters such as the temperature decrease, the turbidity increase, and changes in the host physiology through the gemmulation, bleaching and degradation periods. Additionally, the differences of environmental parameters between sites might not be contrasted enough to generate clear morphological and physiological differences within sponge hosts.

### Temporal changes of the bacterial microbiome through the sexual stage and the senescence of *S. lacustris*

Integrating NMDS, db-RDA, variance partitioning, and morphological observations revealed that the primary drivers of microbial community dynamics were temporal changes in both physico-chemical water conditions and host physiology. Notably, temporal variations have previously been identified as the dominant factor shaping the bacterial communities of *S. lacustris* juveniles originating from the same sponge population sampled at site S3 in this study (Paix *et al*., 2024). In that earlier work, microbial dynamics were linked to successive development stages during early, with community shifts occurring during gemmule hatching followed by the development of choanocytes, the aquiferous system and ultimately the osculum. Prior to these stages, vertical transmission of bacterial symbionts from maternal tissues to the interior of gemmules was suggested, particularly involving members of the *Terasakiellaceae*, which may be associated with thesocytes during gemmulation. In the present study, *Terasakiellaceae* were also consistently detected, notably as a dominant core family present throughout all sampling times across all three sites. The relative abundance of this family decreased over time following gemmulation, reaching lower abundance at T6, when sponges were fully bleached. Taken together with previous evidence for vertical transmission, these findings support the essential role of *Terasakiellaceae* as core endosymbiotic microbial members actively associated with sponge function. Moreover, *Terasakiellaceae* were detected exclusively in freshwater sponge samples and were absent from surrounding water samples, corroborating previous observations from western North Carolina (U.S.A.) across three other freshwater sponge species (Keleher *et al*., 2025).

Additionally, temporal changes in sponge-associated bacterial communities were characterized by an increase in β-diversity dispersion from T4 to T6, while freshwater samples displayed higher homogeneity among replicates. Consequently, increasing stochastic processes in community assembly may explain the greater heterogeneity observed among sponge microbiome replicates. For instance, sponge senescence associated with bleaching may lead to reduced host control over the microbiome, allowing the proliferation of opportunistic bacteria capable of degrading host tissue or utilizing released organic matter. This hypothesis is consistent with the Anna Karenina Principle (AKP), which proposes that increasing β-diversity dispersion can result from dysbiosis induced by host-associated stressors (Zaneveld *et al*., 2017). Similar patterns were previously reported for the *S. lacustris* juveniles subjected to starvation stress, where increased heterogeneity among replicates was observed (Paix *et al*., 2024). In the present study, a different type of stress may likewise lead to a comparable scenario, promoting stochastic assembly of a novel microbial community adapted to host senescence. Such stochastic processes could arise from the loss of essential symbionts involved in maintaining microbiome stability. These “keystone taxa” are thought to play a critical role in preserving microbial community balance and holobiont functioning as demonstrated in other host-microbiome systems (Banerjee *et al*., 2018). In marine sponges, photosymbionts can fulfil such stability role, as shown Candidatus *Synechococcus feldmannii* in *Petrosia ficiformis* (Britstein *et al*., 2020). In this study, *Chlorella*-like photosymbionts might similarly act as “keystone taxa”, promoting determinist processes in microbial community assembly. Their decline during bleaching, linked to decreasing temperature and gemmulation, may therefore trigger a dysbiosis state, facilitating the emergence of a new microbial niche within senescent tissues, driven by stochastic colonization. Differential abundance analyses based on bleaching stage revealed a wide diversity of biomarker taxa across multiple phyla associated with fully bleached sponges. This high diversity is unlikely to reflect pathogens taxa only, but rather suggests colonization by opportunistic microbes scavenging organic matter released during host senescence. To further elucidate these processes, future studies should focus on the diversity of sponge associated microalgal communities, for example through metabarcoding of the ITS, *rbcL* or 23S rRNA gene markers for example, as well as on functional approaches. In particular, characterization of the freshwater sponge holometabolome could provide key insights into physiological changes and stress responses of the holobiont, such as oxidative stress. Moreover, metagenomics and metatranscriptomics approaches would allow more detailed investigation of the functional potential and activity of putative pathogens and microbial scavengers.

## Conclusion

Using an *in-situ* approach, this study revealed the relative importance of temporal and spatial factors shaping the bacterial microbiome of *Spongilla lacustris*. Among these, changes in host physiology, together with decreases in temperature and increase in turbidity, emerged as the primary drivers. As adult sponges transitioned from green, healthy and asexually reproducing individuals to bleached and senescent states, their microbiomes exhibited pronounced shifts in β-diversity, accompanied by increased dispersion consistent with a dysbiosis process. In parallel, spatial differences associated with contrasting nutrient levels revealed enrichment of denitrifying and methanotrophic taxa, highlighting the influence of eutrophication on specific microbial groups. Despite these variations, a conserved core community (e.g. the *Terasakiellaceae*), persisted across all sites and sampling times, underscoring strong and stable associations with freshwater sponges. Together, these initial insights into freshwater sponge-microbiome dynamics emphasize the need for further investigation into functional responses to environmental stressors in urbanized freshwater ecosystems.

## Declarations

### Availability of data and material

16S rRNA gene sequences for metabarcoding were deposited and are publicly available in the NCBI Sequences Read Archive (SRA) under the BioProject ID PRJNA1371752. The R script used for the 16S rRNA gene metabarcoding analyses can be found at https://github.com/Chloen-Lhm/Spongilla-lacustris-microbiome-analyses.

### Competing interests

The authors declare that they have no competing interests.

### Funding

This work was funded by the NWO-VIDI with project number 16.161.301.

### Authors’ contributions

BP and NdV designed the study and performed the fieldwork. BP and CL performed the lab work. CL processed the data. CL and BP analyzed the results. CL, BP, and NdV wrote the manuscript.

## Acknowledgments

We thank Mike Hynes, Anouk Langerak, Jerry Weidema, Maarten Teske, and Joost de Bruijn for their help during the sampling, and Niels van der Windt for his help in the lab. We thank Jan-Willem Versluijs from Jachthaven Lockhorst for his hospitality and for granting us access to his premises for sponge sampling.

## Notes

### Competing Interest Statement

The authors have declared no competing interest.

## References

Aoki, M., Takemura, Y., Kawakami, S., Yoochatchaval, W., Tran P., T., Tomioka, N., et al. (2023) Quantitative detection and reduction of potentially pathogenic bacterial groups of Aeromonas, Arcobacter, Klebsiella pneumoniae species complex, and Mycobacterium in wastewater treatment facilities. PLoS ONE 18: e0291742.

Banerjee, S., Schlaeppi, K., and van der Heijden, M.G.A. (2018) Keystone taxa as drivers of microbiome structure and functioning. Nat Rev Microbiol 16: 567–576.

Baquiran, J.I.P. and Conaco, C. (2018) Sponge-microbe partnerships are stable under eutrophication pressure from mariculture. Marine Pollution Bulletin 136: 125–134.

Belikov, S., Belkova, N., Butina, T., Chernogor, L., Kley, A.M.-V., Nalian, A., et al. (2019) Diversity and shifts of the bacterial community associated with Baikal sponge mass mortalities. PLOS ONE 14: e0213926.

Bennett, H.M., Altenrath, C., Woods, L., Davy, S.K., Webster, N.S., and Bell, J.J. (2017) Interactive effects of temperature and *P* CO _2_ on sponges: from the cradle to the grave. Global Change Biology 23: 2031–2046.

Borrel, G., Jézéquel, D., Biderre-Petit, C., Morel-Desrosiers, N., Morel, J.-P., Peyret, P., et al. (2011) Production and consumption of methane in freshwater lake ecosystems. Research in Microbiology 162: 832–847.

Britstein, M., Cerrano, C., Burgsdorf, I., Zoccarato, L., Kenny, N.J., Riesgo, A., et al. (2020) Sponge microbiome stability during environmental acquisition of highly specific photosymbionts. Environmental Microbiology 22: 3593–3607.

Callahan, B.J., McMurdie, P.J., and Holmes, S.P. (2017) Exact sequence variants should replace operational taxonomic units in marker-gene data analysis. ISME J 11: 2639–2643.

Callahan, B.J., McMurdie, P.J., Rosen, M.J., Han, A.W., Johnson, A.J.A., and Holmes, S.P. (2016) DADA2: High-resolution sample inference from Illumina amplicon data. Nat Methods 13: 581–583.

Carey, R.O. and Migliaccio, K.W. (2009) Contribution of Wastewater Treatment Plant Effluents to Nutrient Dynamics in Aquatic Systems: A Review. Environmental Management 44: 205–217.

Dahal, R.H., Chaudhary, D.K., and Kim, J. (2017) Rurimicrobium arvi gen. nov., sp. nov., a member of the family Chitinophagaceae isolated from farmland soil. International Journal of Systematic and Evolutionary Microbiology 67: 5235–5243.

Davis, N.M., Proctor, D.M., Holmes, S.P., Relman, D.A., and Callahan, B.J. (2018) Simple statistical identification and removal of contaminant sequences in marker-gene and metagenomics data. Microbiome 6: 226.

De Castro-Fernández, P., Ballesté, E., Angulo-Preckler, C., Biggs, J., Avila, C., and García-Aljaro, C. (2023) How does heat stress affect sponge microbiomes? Structure and resilience of microbial communities of marine sponges from different habitats. Frontiers in Marine Science 9:.

Emmanuel, A., Wei, Y., Ramzan, M.N., Yang, W., and Zheng, Z. (2025) Dynamics of Bacterial Communities and Their Relationship with Nutrients in a Full-Scale Shrimp Recirculating Aquaculture System in Brackish Water. Animals 15: 1400.

Foster, Z.S.L., Sharpton, T.J., and Grünwald, N.J. (2017) Metacoder: An R package for visualization and manipulation of community taxonomic diversity data. PLOS Computational Biology 13: e1005404.

Gantt, S.E., McMurray, S.E., Stubler, A.D., Finelli, C.M., Pawlik, J.R., and Erwin, P.M. (2019) Testing the relationship between microbiome composition and flux of carbon and nutrients in Caribbean coral reef sponges. Microbiome 7: 124.

Gernert, C., Glöckner, F.O., Krohne, G., and Hentschel, U. (2005) Microbial diversity of the freshwater sponge *Spongilla lacustris*. Microb Ecol 50: 206–212.

Gilbert, J.J. (1975) Field Experiments on Gemmulation in the Fresh-Water Sponge Spongilla lacustris. Transactions of the American Microscopical Society 94: 347.

Glasl, B., Luter, H.M., Damjanovic, K., Kitzinger, K., Mueller, A.J., Mahler, L., et al. (2024) Co-occurring nitrifying symbiont lineages are vertically inherited and widespread in marine sponges. The ISME Journal wrae069.

Gloor, G.B., Macklaim, J.M., Pawlowsky-Glahn, V., and Egozcue, J.J. (2017) Microbiome datasets are compositional: and this is not optional. Front Microbiol 8: 2224.

Graffius, S., Garzón, J.F.G., Zehl, M., Pjevac, P., Kirkegaard, R., Flieder, M., et al. (2023) Secondary metabolite production potential in a microbiome of the freshwater sponge *Spongilla lacustris*. Microbiology Spectrum 11: e04353–22.

Hall, C., Camilli, S., Dwaah, H., Kornegay, B., Lacy, C., Hill, M.S., and Hill, A.L. (2021) Freshwater sponge hosts and their green algae symbionts: a tractable model to understand intracellular symbiosis. PeerJ 9: e10654.

Han, B.-N., Hong, L.-L., Gu, B.-B., Sun, Y.-T., Wang, J., Liu, J.-T., and Lin, H.-W. (2019) Natural Products from Sponges. In Symbiotic Microbiomes of Coral Reefs Sponges and Corals. Li, Z. (ed). Dordrecht: Springer Netherlands, pp. 329–463.

Jensen, K.S.- and Pedersen, M.F. (1994) Photosynthesis by symbiotic algae in the freshwater sponge, Spongilla lacustris. Limnol Oceanogr 39: 551–561.

Kasozi, N., Kaiser, H., and Wilhelmi, B. (2020) Metabarcoding Analysis of Bacterial Communities Associated with Media Grow Bed Zones in an Aquaponic System. International Journal of Microbiology 2020: 8884070.

Keleher, J.G., Strope, T.A., Estrada, N.E., Griggs Mathis, A.M., Easson, C.G., and Fiore, C. (2025) Freshwater sponges in the southeastern U.S. harbor unique microbiomes that are influenced by host and environmental factors. PeerJ 13: e18807.

Klindworth, A., Pruesse, E., Schweer, T., Peplies, J., Quast, C., Horn, M., and Glöckner, F.O. (2013) Evaluation of general 16S ribosomal RNA gene PCR primers for classical and next-generation sequencing-based diversity studies. Nucleic Acids Res 41: e1.

Kulakova, N.V., Sakirko, M.V., Adelshin, R.V., Khanaev, I.V., Nebesnykh, I.A., and Pérez, T. (2018) Brown rot syndrome and changes in the bacterial сommunity of the Baikal sponge *Lubomirskia baikalensis*. Microb Ecol 75: 1024–1034.

Lahti, L., Shetty, S., Blake, T., and Salojarvi, J. (2017) Tools for microbiome analysis in R. Version 1: 28.

Leys, S., Grombacher, L., and Hill, A. (2019) Hatching and freezing gemmules from the freshwater sponge Ephydatia muelleri.

Lo Giudice, A. and Rizzo, C. (2024) Freshwater Sponges as a Neglected Reservoir of Bacterial Biodiversity. Microorganisms 12: 25.

Manconi, R. and Pronzato, R. (2008) Global diversity of sponges (Porifera: Spongillina) in freshwater. In Freshwater Animal Diversity Assessment. Developments in Hydrobiology. Balian, E.V., Lévêque, C., Segers, H., and Martens, K. (eds). Dordrecht: Springer Netherlands, pp. 27–33.

Martinez Arbizu, P. (2020) pairwiseAdonis: Pairwise multilevel comparison using adonis. R package version 04 1:.

Maslin, M., Paix, B., van der Windt, N., Ambo-Rappe, R., Debitus, C., Gaertner-Mazouni, N., et al. (2024) Prokaryotic communities of the French Polynesian sponge *Dactylospongia metachromia* display a site-specific and stable diversity during an aquaculture trial. Antonie van Leeuwenhoek 117: 65.

McMurdie, P.J. and Holmes, S. (2013) phyloseq: an R package for reproducible interactive analysis and graphics of microbiome census data. PLoS One 8: e61217.

McMurdie, P.J. and Holmes, S. (2014) Waste not, want not: why rarefying microbiome data is inadmissible. PLoS Comput Biol 10: e1003531.

Oksanen, J., Blanchet, F.G., Friendly, M., Kindt, R., Legendre, P., McGlinn, D., et al. (2019) vegan: Community ecology package.

Osinga, R., Armstrong, E., Grant Burgess, J., Hoffmann, F., Reitner, J., and Schumann-Kindel, G. (2001) Sponge–microbe associations and their importance for sponge bioprocess engineering. Hydrobiologia 461: 55–62.

Paix, B., van der Valk, E., and de Voogd, N.J. (2024) Dynamics, diversity, and roles of bacterial transmission modes during the first asexual life stages of the freshwater sponge Spongilla lacustris. Environmental Microbiome 19: 37.

Pelsma, K.A.J., Verhagen, D.A.M., Dean, J.F., Jetten, M.S.M., and Welte, C.U. (2023) Methanotrophic potential of Dutch canal wall biofilms is driven by *Methylomonadaceae*. FEMS Microbiology Ecology 99: fiad110.

Pita, L., Rix, L., Slaby, B.M., Franke, A., and Hentschel, U. (2018) The sponge holobiont in a changing ocean: from microbes to ecosystems. Microbiome 6: 46.

Posadas, N., Baquiran, J.I.P., Nada, M.A.L., Kelly, M., and Conaco, C. (2022) Microbiome diversity and host immune functions influence survivorship of sponge holobionts under future ocean conditions. ISME J 16: 58–67.

Pronzato, R., Pisera, A., and Manconi, R. (2017) Fossil freshwater sponges: taxonomy, geographic distribution, and critical review. Acta Palaeontologica Polonica 62: 468–495.

Qian, X., Huang, J., Cao, C., Yao, J., Wu, Y., Wang, L., and Wang, X. (2024) Modified basalt fiber filled in constructed wetland-microbial fuel cell: Comparison of performance and microbial impacts under PFASs exposure. Journal of Hazardous Materials 476: 135179.

Quast, C., Pruesse, E., Yilmaz, P., Gerken, J., Schweer, T., Yarza, P., et al. (2013) The SILVA ribosomal RNA gene database project: improved data processing and web-based tools. Nucleic Acids Res 41: D590–D596.

Radax, R., Hoffmann, F., Rapp, H.T., Leininger, S., and Schleper, C. (2012) Ammonia-oxidizing archaea as main drivers of nitrification in cold-water sponges. Environ Microbiol 14: 909–923.

Regoli, F., Cerrano, C., Chierici, E., Bompadre, S., and Bavestrello, G. (2000) Susceptibility to oxidative stress of the Mediterranean demosponge Petrosia ficiformisL: role of endosymbionts and solar irradiance. Marine Biology 137: 453–461.

Ribeiro, S.M., Omena, E.P., and Muricy, G. (2003) Macrofauna associated to *Mycale microsigmatosa* (Porifera, Demospongiae) in Rio de Janeiro State, SE Brazil. Estuarine, Coastal and Shelf Science 57: 951–959.

Ribes, M., Calvo, E., Movilla, J., Logares, R., Coma, R., and Pelejero, C. (2016) Restructuring of the sponge microbiome favors tolerance to ocean acidification. Environmental Microbiology Reports 8: 536–544.

Ribes, M., Yahel, G., Romera-Castillo, C., Mallenco, R., Morganti, T.M., and Coma, R. (2023) The removal of dissolved organic matter by marine sponges is a function of its composition and concentration: An in situ seasonal study of four Mediterranean species. Science of The Total Environment 871: 161991.

Rix, L., de Goeij, J.M., van Oevelen, D., Struck, U., Al-Horani, F.A., Wild, C., and Naumann, M.S. (2017) Differential recycling of coral and algal dissolved organic matter via the sponge loop. Functional Ecology 31: 778–789.

Rizzo, C., Caruso, G., Maimone, G., Patrolecco, L., Termine, M., Bertolino, M., et al. (2025) Microbiome and pollutants in the freshwater sponges Ephydatia muelleri (Lieberkühn, 1856) and Spongilla lacustris (Linnaeus, 1758) from the sub-Arctic Pasvik river (Northern Fennoscandia). Environmental Research 273: 121126.

Ruprecht, J.E., Birrer, S.C., Dafforn, K.A., Mitrovic, S.M., Crane, S.L., Johnston, E.L., et al. (2021) Wastewater effluents cause microbial community shifts and change trophic status. Water Research 200: 117206.

Rützler, K. (2012) The Role of Sponges in the Mesoamerican Barrier-Reef Ecosystem, Belize. In Advances in Marine Biology. Elsevier, pp. 211–271.

Simister, R., Taylor, M.W., Tsai, P., and Webster, N. (2012) Sponge-microbe associations survive high nutrients and temperatures. PLoS One 7: e52220.

Thomas, T., Moitinho-Silva, L., Lurgi, M., Björk, J.R., Easson, C., Astudillo-García, C., et al. (2016) Diversity, structure and convergent evolution of the global sponge microbiome. Nat Commun 7: 11870.

Tian, R., Sun, J., Cai, L., Zhang, W., Zhou, G., Qiu, J., and Qian, P. (2016) The deepLsea glass sponge *Lophophysema eversa* harbours potential symbionts responsible for the nutrient conversions of carbon, nitrogen and sulfur. Environmental Microbiology 18: 2481–2494.

Turner, C.R., Miller, D.J., Coyne, K.J., and Corush, J. (2014) Improved methods for capture, extraction, and quantitative assay of environmental DNA from asian bigheaded carp (*Hypophthalmichthys* spp.). PLOS ONE 9: e114329.

Wang, H., Lin, L., Zhang, L., Han, P., and Ju, F. (2023) Microbiome assembly mechanism and functional potential in enhanced biological phosphorus removal system enriched with Tetrasphaera-related polyphosphate accumulating organisms. Environmental Research 233: 116494.

Webster, N.S. and Thomas, T. (2016) The Sponge Hologenome. mBio 7: e00135–00116.

Wilkinson, C.R. (1980) Nutrient translocation from green algal symbionts to the freshwater sponge *Ephydatia fluviatilis*. Hydrobiologia 75: 241–250.

Williamson, C.E. (1979) An Ultrastructural Investigation of Algal Symbiosis in White and Green Spongilla lacustris (L.) (Porifera: Spongillidae). Transactions of the American Microscopical Society 98: 59–77.

Zaneveld, J.R., McMinds, R., and Vega Thurber, R. (2017) Stress and stability: applying the Anna Karenina principle to animal microbiomes. Nat Microbiol 2: 1–8.

