## Supplementary information for "Host senescence and urbanized conditions drive bacterial communities of the freshwater sponge *Spongilla lacustris*"

### Supplementary tables

#### Table S1. Summary of the sample list describing the sampling times, dates, sites, number of replicates and the bleaching stage of the sponge replicates. ^a^ No samples were collected on T1 for S1 and S2, as *S. lacustris* populations in these sites were only observed since T2.

| **Sampling**  **times** ^a^ | **Sampling sites** | **Sampling date** | **Number of sponge sample replicates** | **Bleaching stage of the sponge tissues** | **Number of freshwater sample replicates** |
| --- | --- | --- | --- | --- | --- |
| T1 | S3 | 01/10/2021 | 4 | Green (all) | 4 |
| T2 | S1 | 14/10/2021 | 4 | Green (all) | 3 |
|  | S2 | 14/10/2021 | 4 | Green (all) | 2 |
|  | S3 | 15/10/2021 | 4 | Green (all) | 3 |
| T3 | S1 | 28/10/2021 | 4 | Green (3) and partially bleached (1) | 4 |
|  | S2 | 29/10/2021 | 4 | Green (3) and fully bleached (1) | 4 |
|  | S3 | 27/10/2021 | 4 | Green (all) | 4 |
| T4 | S1 | 12/11/2021 | 4 | Partially bleached (3) and fully bleached (1) | 4 |
|  | S2 | 11/11/2021 | 4 | Partially bleached (all) | 4 |
|  | S3 | 10/11/2021 | 4 | Green (2), fully bleached (1), partially bleached (1) | 4 |
| T5 | S1 | 24/11/2021 | 4 | Partially bleached (1) , fully bleached (3) | 4 |
|  | S2 | 25/11/2021 | 4 | Green (1), partially bleached (2), fully bleached (1) | 4 |
|  | S3 | 23/11/2021 | 4 | Green (1), partially bleached (3) | 4 |
| T6 | S1 | 14/12/2021 | 4 | Partially bleached (1), fully bleached (3) | 4 |
|  | S2 | 15/12/2021 | 3 | Fully bleached (all) | 3 |
|  | S3 | 13/12/2021 | 4 | Fully bleached (all) | 4 |

#### Table S2. Results of the Shapiro test for α-diversity indexes (Chao1, Pielou and Shannon).

|  | **All samples** | | **Sponge samples only** | |
| --- | --- | --- | --- | --- |
|  | **W** | ***p*** | **W** | ***p*** |
| **Chao1** | 0.93335 | 1.976e-05 | 0.92175 | **0.000728** |
| **Pielou** | 0.93421 | 2.242e-05 | 0.95018 | **0.01362** |
| **Shannon** | 0.96918 | 0.008552 | 0.96569 | 0.08035 |

#### Table S3. Results from Kruskal-Wallis tests and two-way ANOVA conducted with the α-diversity indices (Chao1, Pielou and Shannon) of sponge samples, with the sampling times and sites as factor of comparisons. D.f., F and *p* correspond to degrees of freedom; F ratio and p-value, respectively.

|  | | **Chi-square** | | **D.f.** | | ***p*** |
| --- | --- | --- | --- | --- | --- | --- |
| **Chao1** | **Times** | 3.408137 | | 5 | | 0.63733 |
|  | **Sites** | 9.8363 | | 2 | | **0.00731** |
| **Pielou** | **Times** | 16.178979 | | 5 | | **0.00635** |
|  | **Sites** | 3.623505 | | 2 | | 0.16337 |
|  | | **D.f.** | **Sum of squares** | **Mean square** | **F** | ***p*** |
| **Shannon** | **Times** | 5 | 0.39 | 0.0785 | 0.144 | 0.9814 |
|  | **Sites** | 2 | 3.19 | 1.5793 | 2.897 | **0.0598** |
|  | **Times*Sites** | 8 | 1.55 | 0.1936 | 0.355 | 0.9414 |
|  | **Residual** | 101 | 55.06 | 0.5452 |  |  |

#### Table S4. Results of the PERMANOVA test of the *β*-diversity conducted with the “Sample type” factor (freshwater *vs* sponges). D.f., F, R^2^ and *p* correspond to degrees of freedom, F ratio, coefficient of determination, and *p*-value, respectively

| **Factor** | **Df** | **R²** | **F** | ***p*** |
| --- | --- | --- | --- | --- |
| **Sample type** | 1 | 0.362 | 65.322 | **0.001** |
| **Residual** | 115 | 16.648 | 0.638 |  |
| **Total** | 116 | 26.104 | 1.000 |  |

#### Table S5. Results of the nested PERMANOVA test of the *β*-diversity of sponge samples, conducted with the “sampling time” and “sampling site” factors. D.f., F, R^2^ and *p* correspond to degrees of freedom, F ratio, coefficient of determination, and *p*-value, respectively.

| **Factor** | **Df** | **R²** | **F** | ***p*** |
| --- | --- | --- | --- | --- |
| **Sampling time** | 5 | 0.341 | 6.792 | **0.001** |
| **Sampling site** | 2 | 0.064 | 3.185 | **0.001** |
| **Sampling site*Sampling time** | 8 | 0.134 | 1.676 | **0.001** |
| **Residual** | 101 | 18.922 | 0.725 |  |
| **Total** | 116 | 26.104 | 1 |  |

#### Table S6. Results of the multivariate pairwise adonis test comparing the *β*-diversity of sponge samples between the sampling times. F, R^2^, and p correspond to the F ratio, coefficient of determination, and *p*-value, respectively

| **Pairwise comparison** | **Sum of squares** | **F** | **R²** | ***p*** |
| --- | --- | --- | --- | --- |
| **T1 vs T2** | 0.008 | 1.871 | 0.118 | 0.125 |
| **T1 vs T3** | 0.031 | 13.226 | 0.486 | 0.002 |
| **T1 vs T4** | 0.032 | 6.964 | 0.332 | 0.002 |
| **T1 vs T5** | 0.030 | 9.147 | 0.413 | 0.002 |
| **T1 vs T6** | 0.055 | 14.678 | 0.530 | 0.002 |
| **T2 vs T3** | 0.020 | 5.608 | 0.203 | 0.002 |
| **T2 vs T4** | 0.026 | 5.303 | 0.194 | 0.001 |
| **T2 vs T5** | 0.036 | 8.617 | 0.291 | 0.001 |
| **T2 vs T6** | 0.146 | 32.961 | 0.611 | 0.001 |
| **T3 vs T4** | 0.025 | 7.1367 | 0.245 | 0.001 |
| **T3 vs T5** | 0.061 | 22.655 | 0.519 | 0.001 |
| **T3 vs T6** | 0.250 | 83.230 | 0.799 | 0.001 |
| **T4 vs T5** | 0.014 | 3.308 | 0.136 | 0.033 |
| **T4 vs T6** | 0.179 | 39.693 | 0.654 | 0.001 |
| **T5 vs T6** | 0.112 | 30.711 | 0.606 | 0.001 |

#### Table S7. Results of the multivariate pairwise adonis test comparing the *β*-diversity of sponge samples between sampling sites. F, R^2^, and p correspond to the F ratio, coefficient of determination, and *p*-value, respectively

| **Pairwise comparison** | **Sum of squares** | **F** | **R²** | ***p*** |
| --- | --- | --- | --- | --- |
| **S3 vs S2** | 0.009 | 1.037 | 0.025 | 0.346 |
| **S3 vs S1** | 0.006 | 0.692 | 0.017 | 0.489 |
| **S2 vs S1** | 0.010 | 1.030 | 0.027 | 0.327 |

#### Table S8. Results from Kruskal-Wallis tests conducted with the *β*-dispersion of sponge samples, with the sampling times as factor of comparisons, for each sampling site. D.f. and p correspond to degrees of freedom, and p-value, respectively.

| **Comparisons** | **Chi-square** | **D.f.** | ***p*** |
| --- | --- | --- | --- |
| **Sampling times within S1** | 12.429 | 4 | **0.014** |
| **Sampling times within S2** | 13.595 | 4 | **0.009** |
| **Sampling times within S3** | 17.505 | 5 | **0.004** |

#### Table S9. Results from the two-way ANOVA test comparing the relative richness of core ASVs of sponges according to the sampling site and sampling times factors. D.f., F and *p* correspond to degrees of freedom; F ratio and p-value, respectively.

|  | **D.f.** | **Sum of squares** | **Mean square** | **F** | ***p*** |
| --- | --- | --- | --- | --- | --- |
| Site | 2 | 81.82 | 40.91 | 7.289 | **0.00178** |
| Date | 5 | 11.29 | 2.26 | 0.402 | 0.84460 |
| Site*Date | 8 | 70.46 | 8.81 | 1.569 | 0.16047 |
| Residuals | 46 | 258.17 | 5.61 |  |  |

### Supplementary figures

#### Figure S1. Photographs of *Spongilla lacustris* specimens collected. A. Photographs of *S. lacustris* for each sampling site and time. B. Photograph of a sample of *S. lacustris* fully bleached with dark spots marked with the red arrows.

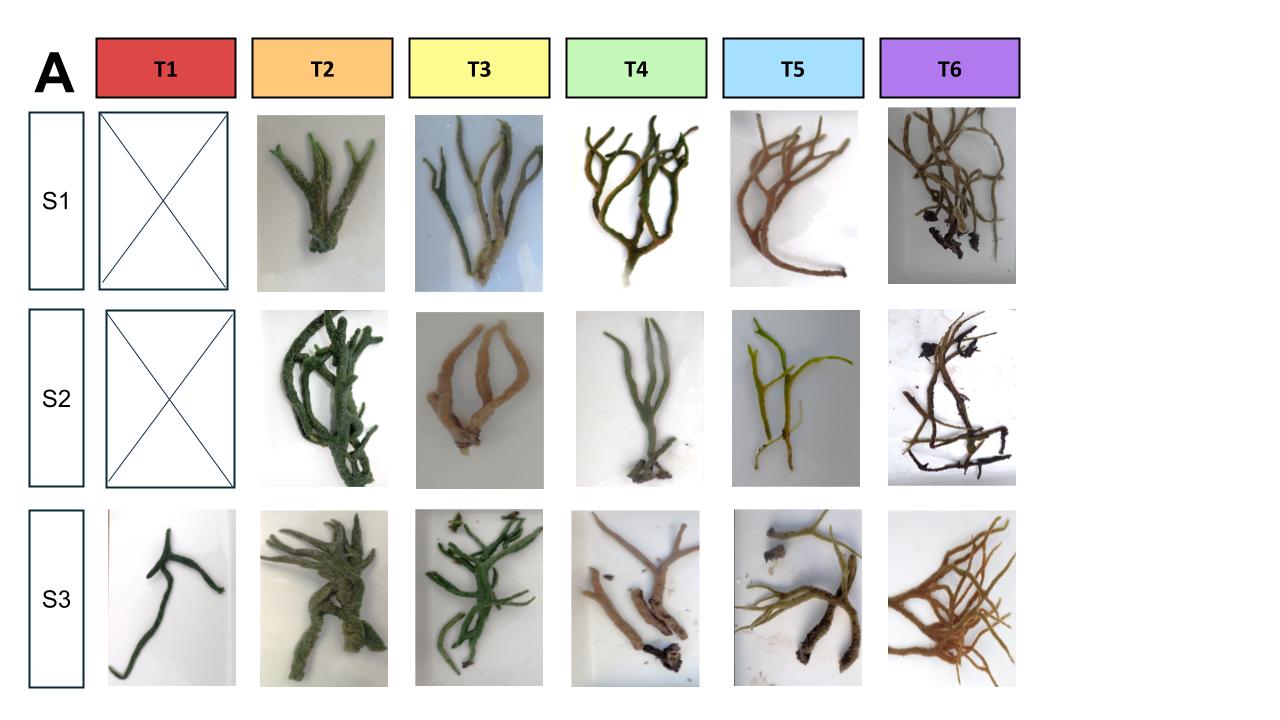

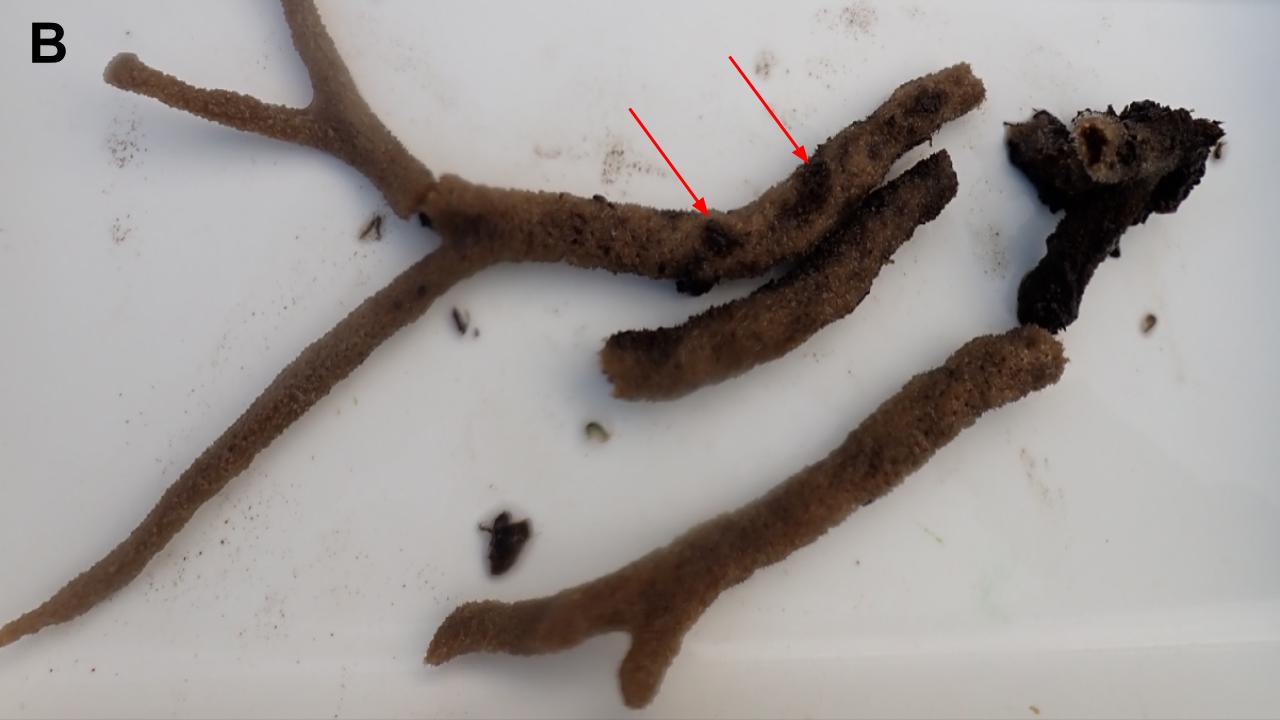

#### Figure S2. Boxplot of the relative weight of the gemmules on the total weight of the sponge (expressed in percentage).

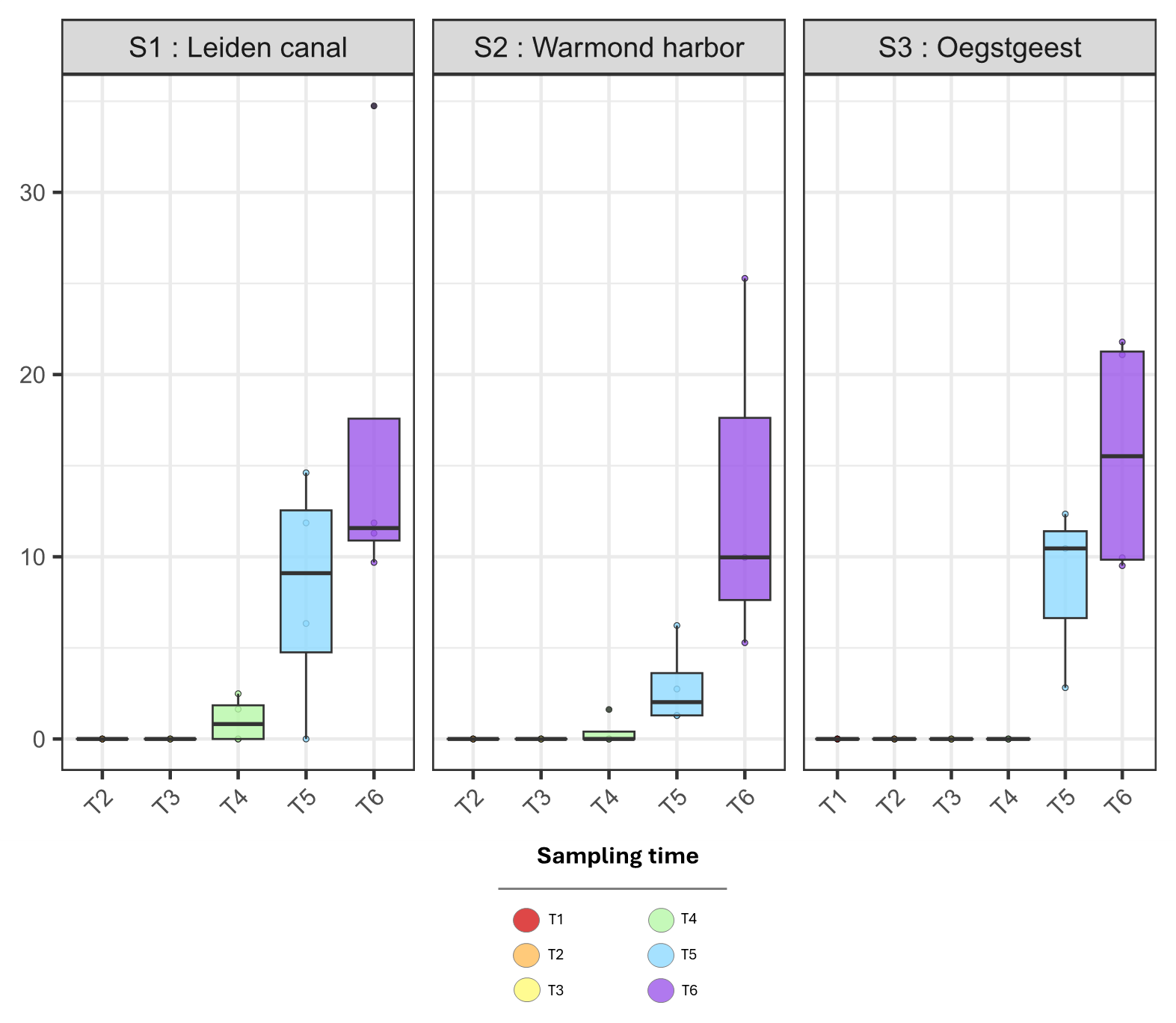

#### Figure S3. Rarefaction curves obtained after the data processing of the 16S rRNA gene sequences with the DADA2 pipeline and the filtration of sequences affiliated to eukaryotes, chloroplasts and mitochondria.

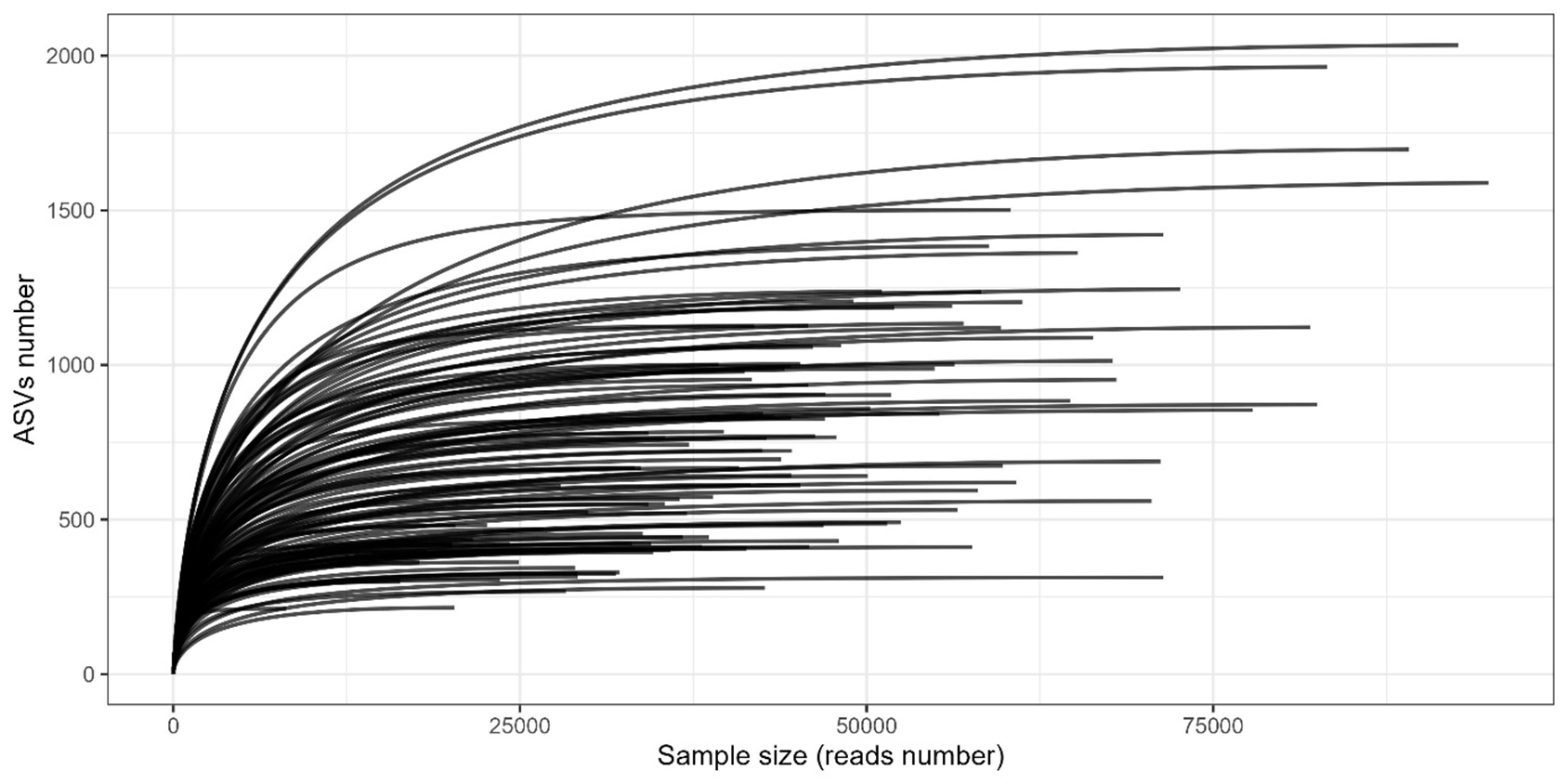

#### Figure S4. Boxplots of the *α*-diversity metrics (Chao1, Shannon and Pielou indices) of the bacterial community of freshwater and *S. lacustris* samples. Lower case indices indicate the results of the Wilcoxon tests for pairwise comparison between sites, and between sampling times.

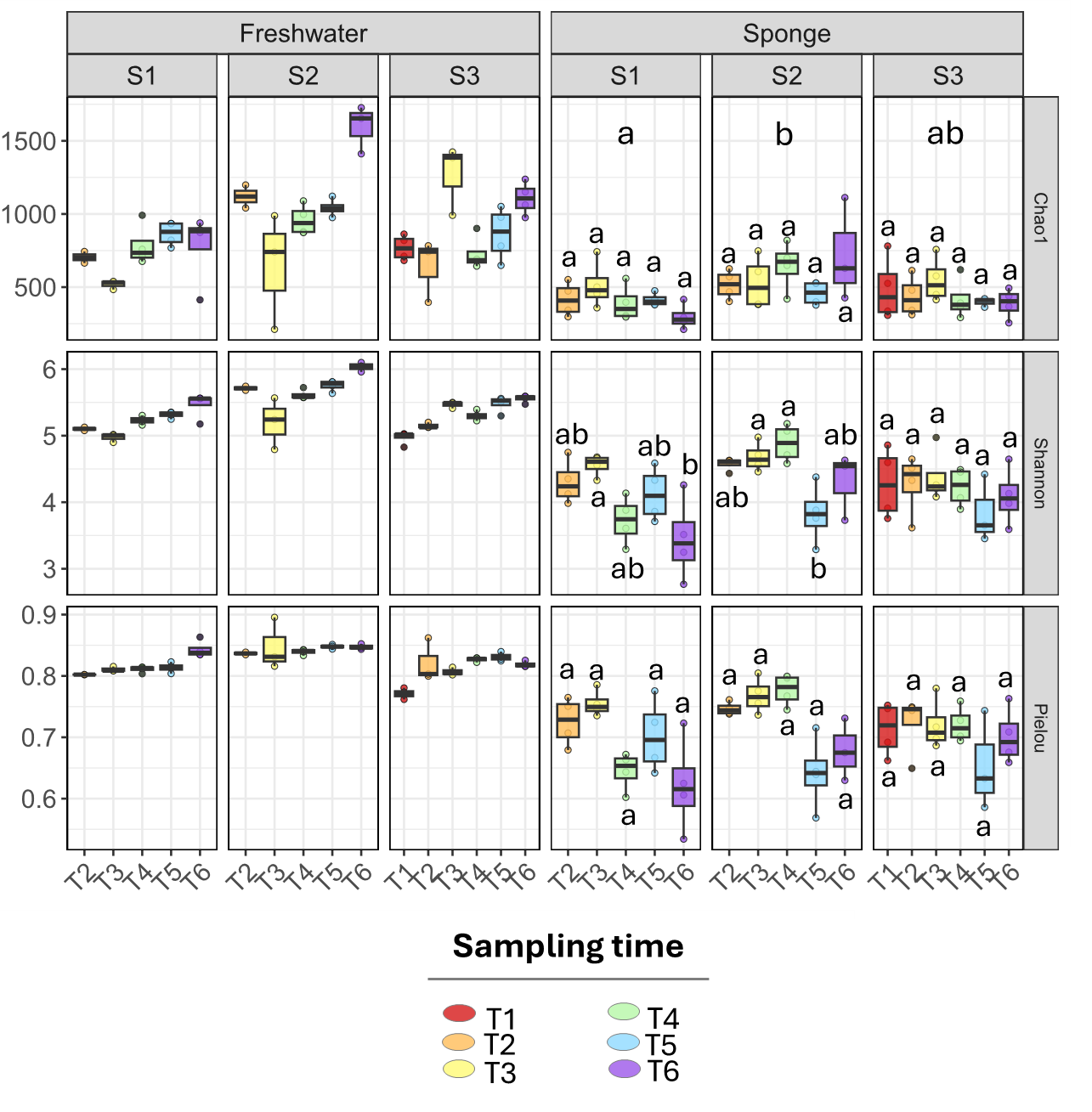

#### Figure S5. Barplots of the bacterial community composition at the family level associated with sponge (A) and freshwater (B) samples. “Other” and “NA” correspond respectively to families with a relative percentage below 5% and unaffiliated families.

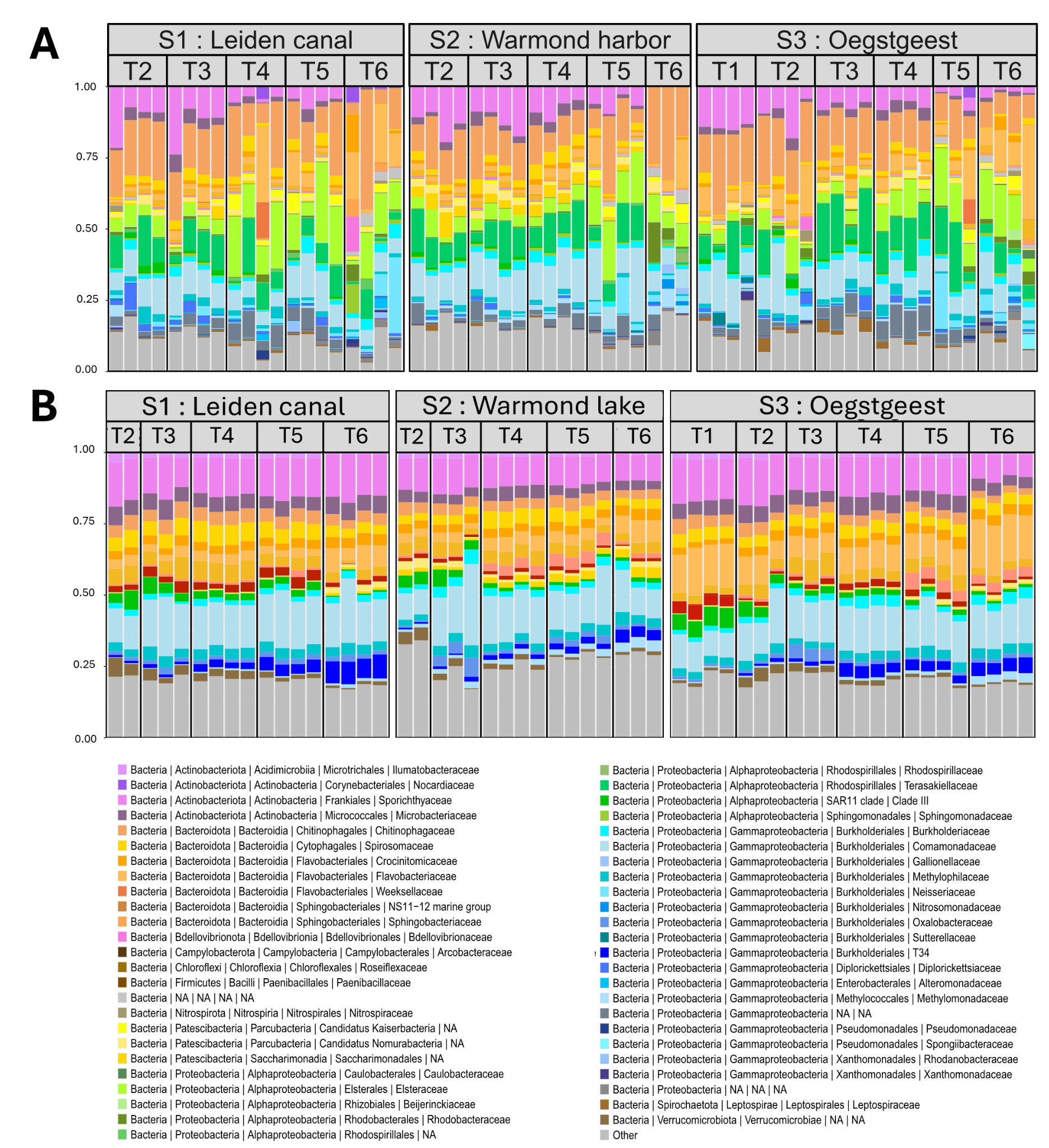

#### Figure S6. Boxplots of the relative core ASVs richness within each sponge sample. Upper- and lower-case indices indicate the results of the HSD Tukey’s tests for pairwise comparison between sites, and between sampling times, respectively.

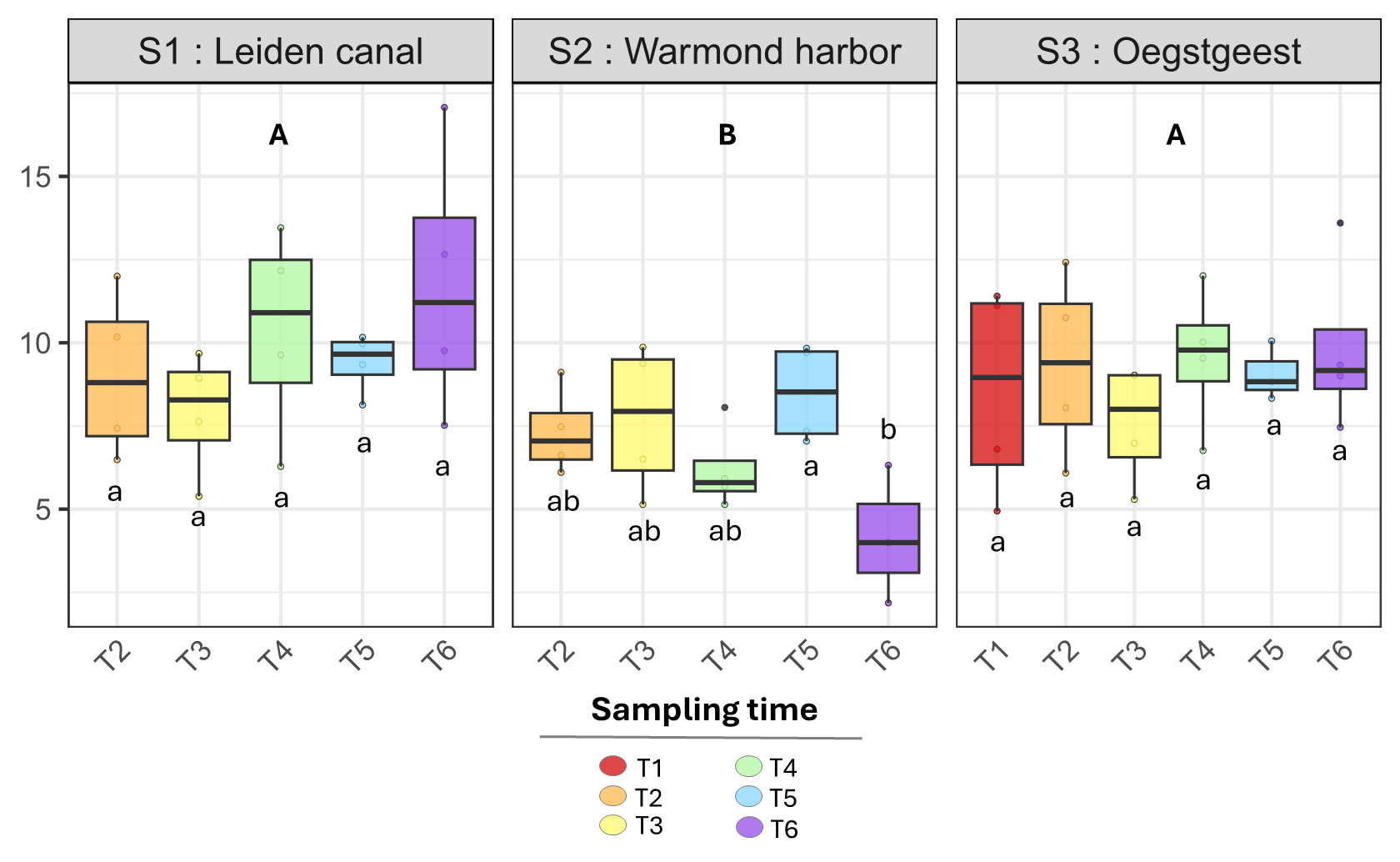

#### Figure S7. Phylogenetic heat trees representing the bacterial taxa significantly and differentially abundant between S2 and S1-S3 (A) and between S3 and S1-S2 (B). For each taxon, (i) the colors of their associated nodes correspond to the log2 fold change between the sites groups, (ii) the size of the nodes corresponds to the relative abundance of each taxon.

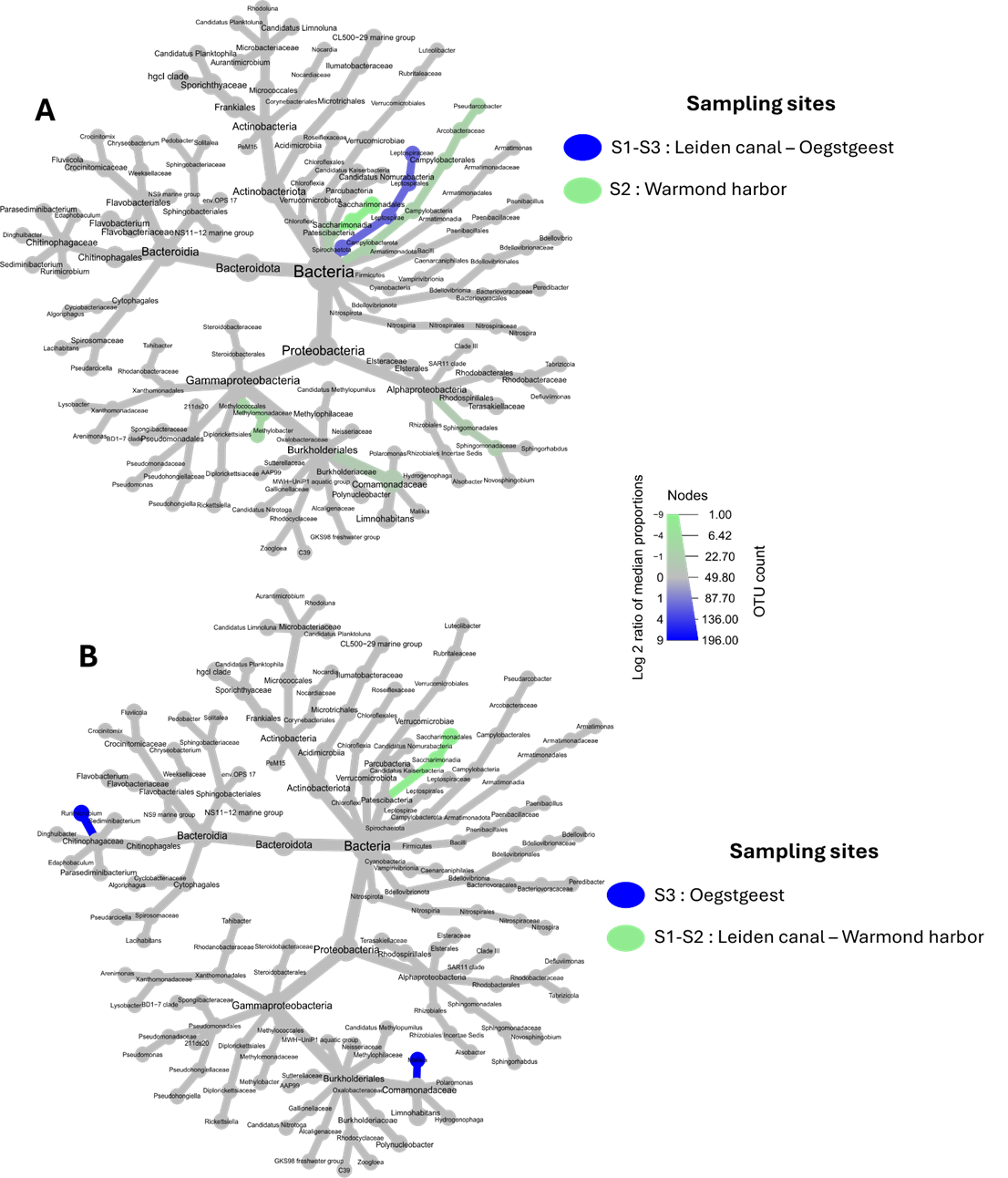
